# Euchromatin factors HULC and Set1C affect heterochromatin organization for mating-type switching in fission yeast *Schizosaccharomyces pombe*

**DOI:** 10.1101/2021.03.23.436714

**Authors:** Alfredo Esquivel-Chávez, Takahisa Maki, Hideo Tsubouchi, Testuya Handa, Hiroshi Kimura, James E. Haber, Genevieve Thon, Hiroshi Iwasaki

## Abstract

Mating-type (P or M) of fission yeast *Schizosaccharomyces pombe* is determined by the transcriptionally active *mat1* cassette and is switched by gene conversion using a donor, either *mat2* or *mat3*, located in an adjacent heterochromatin region (mating-type switching; MTS). In the process, heterochromatic donors of genetic information are selected based on the P or M cell type and on the action of two recombination enhancers, *SRE2* promoting the use of *mat2-P* and *SRE3* promoting the use of *mat3-M*, leading to replacement of the content of the expressed *mat1* cassette. Recently, we found that the histone H3K4 methyltransferase complex Set1C participates in donor selection, raising the question of how a complex best known for its effects in euchromatin controls recombination in heterochromatin. Here, we report that the histone H2BK119 ubiquitin ligase complex HULC functions with Set1C in MTS, as mutants in the *shf1, brl1, brl2* and *rad6* genes showed defects similar to Set1C mutants and belonged to the same epistasis group as *set1*Δ. Moreover, using H3K4R and H2BK119R histone mutants and a Set1-Y897A catalytic mutant indicated that ubiquitylation of histone H2BK119 by HULC and methylation of histone H3K4 by Set1C are functionally coupled in MTS. Cell-type biases in MTS in these mutants suggested that HULC and Set1C inhibit the use of the *SRE3* recombination enhancer in M cells, thus favoring *SRE2* and *mat2-P*. Consistently, imbalanced switching in the mutants was traced to compromised association of the directionality factor Swi6 with the recombination enhancers in M cells. Based on their known effects at other chromosomal locations, we speculate that HULC and Set1C might control nucleosome mobility and strand invasion near the *SRE* elements. In addition, we uncovered distinct effects of HULC and Set1C on histone H3K9 methylation and gene silencing, consistent with additional functions in the heterochromatic domain.

## Introduction

Homothallic strains (*h*^*90*^) of the fission yeast *Schizosaccharomyces pombe* switch between two mating types, P and M. This process is known as mating-type switching (MTS) and takes place at the *mat* locus on chromosome 2. The *mat* locus comprises three cassettes, *mat1, mat2* and *mat3* (Fig. 1A). The active *mat1* cassette expresses either P or M mating-type specific genes and determines the mating-type of a haploid cell (Kelly et al. 1988). The genes in the *mat2* and *mat3* cassettes, P (*mat2-P*) and M (*mat3-M*), respectively, are invariant, and are silenced by heterochromatin. Each cassette is flanked by short homology boxes called *H1* and *H2*. MTS is a result of gene conversion repair of a programmed double-strand break at the *mat1* cassette, using the homology boxes at either the *mat2-P* or *mat3-M* donor cassette (Klar et al. 2014).

**Fig 1.**
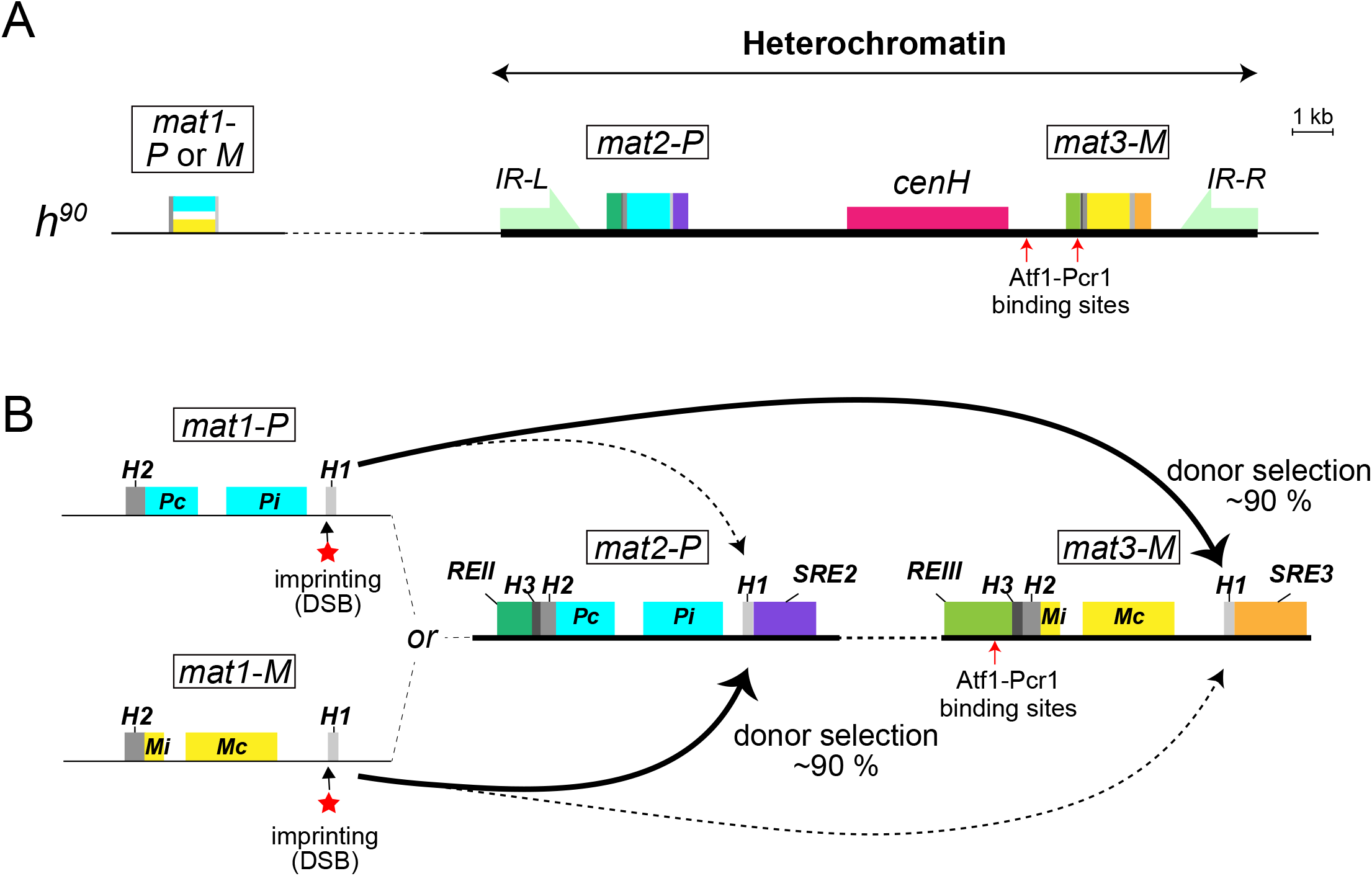
Mating-type switching in *S. pombe*. (A) Schematic representation of the mating-type region. The expressed *mat1* cassette containing either P (blue) or M (yellow) mating-type genes and the silenced donor cassettes, *mat2-P* and *mat3-M* are located in a heterochromatic region bordered by the *IR-L* and *IR-R* repeats. (B) Donor preference of mating-type switching. Each *mat* cassette is flanked by the homology boxes *H2* and *H1, mat2-P* and *mat3-M* have in addition a *H3* box. An imprinting site converted to a single-ended double-strand break (DSB) during DNA replication is located at the junction of *mat1* and its *H1* homology box (depicted here as a red star). Donor preference is determined by the information at *mat1*: *mat1-P* cells select *mat3-M* (∼90%) and *mat1-M* cells select *mat2-P* (∼90%). Red arrows show Atf1-Pcr1 heterodimer binding sites. *REII*: Repressor Element II; *REIII*: Repressor Element III; *Pc*: the P-specific transcription factor Pc gene; *Mc*: the M-specific transcription factor Mc gene; *Pi*: the P-specific transcription factor Pi gene; *Mi*: the M-specific polypeptide Mi gene.

A cell-type specific regulation takes place at the donor-selection step, where cells usually select the mating-type donor cassette opposite to the allele present at *mat1*, thus, P cells (*mat1-P*) preferentially choose *mat3-M* as a donor, while M cells (*mat1-M*) preferentially choose *mat2-P* (Fig. 1B) (Thon et al. 2018). This donor selection, also known as directionality of MTS, requires the mating-type switching factors Swi2 and Swi5, the heterochromatin protein 1 (HP1) homologue Swi6, and two *cis-*acting DNA elements (Egel et al. 1984; Jakociunas et al. 2013; Jia et al. 2004b). The *cis-*acting DNA elements are called <u>S</u>wi2-dependent <u>R</u>ecombination <u>E</u>nhancers, *SRE2* and *SRE3*, and are located next to the *H1* box of *mat2-P* and *mat3-M*, respectively (Jakociunas et al. 2013; Jia et al. 2004b). The Swi2 protein localizes to these elements according to cell type: in P cells, Swi2 localizes only to *SRE3*; in M cells, Swi2 localizes to both *SRE2* and *SRE3* (Jakociunas et al. 2013; Jia et al. 2004b). From yeast two-hybrid assays, Swi2 has been shown to interact with Swi5 and with the recombination factor Rad51; therefore, it has been suggested that the Swi2-Swi5 complex promotes homologous recombination at the *SRE2-* or *SRE3*-adjacent cassette (Akamatsu et al. 2003). Furthermore, the Swi6 protein spreads over the *mat2* and *mat3* region between the flanking boundary elements *IR-L* and *IR-R* (Fig. 1A) and it contributes to form heterochromatin over this entire region (Noma et al. 2001; Thon et al. 2002). The association of Swi6 with the region differs between P and M cells, with greater enrichment of Swi6 in M cells than P cells, although overall cellular levels of Swi6 are comparable between P and M cells (Noma et al. 2001). As Swi2 interacts with Swi6 *in vitro*, the localization pattern of Swi2 is believed to be connected to its interaction with Swi6 (Akamatsu et al. 2003; Akamatsu et al. 2007; Jia et al. 2004b).

Swi6 localization at the *mat* locus is controlled by several histone modifications (Allshire and Madhani 2018). An essential histone modification for Swi6 localization is di- or tri-methylation of histone H3 at lysine 9 (H3K9me2 and -me3, respectively) catalyzed by the methyltransferase Clr4, the homologue of human SUV39H1 and SUV39H2 (Ekwall and Ruusala 1994; Nakayama et al. 2001; Rea et al. 2000). H3K9me2 and -me3 are detected in constitutive heterochromatin including *mat*, centromeres and telomeres, and in facultative heterochromatin such as meiotic genes (Allshire and Ekwall 2015; Zofall et al. 2012).

Constitutive heterochromatic regions contain repetitive DNA sequences that nucleate H3K9 methylation through RNA interference (RNAi) (Allshire and Ekwall 2015). At the *mat* locus, RNAi is triggered by the transcription of *cenH*, homologous to centromeric repeats (Hall et al. 2002; Volpe et al. 2002). dsRNAs originating from *cenH* are cleaved by the ribonuclease Dcr1, and the siRNA products are bound by the RNAi-induced transcriptional silencing complex (RITS) (Buker et al. 2007; Irvine et al. 2006). RITS loaded with siRNAs recruits the Clr4-Rik1-Cul4 complex (CLRC) to methylate H3K9 (Zhang et al. 2008). At the *mat* locus, there is an additional silencing pathway involving the CREB-like transcription factor, Atf1-Pcr1 (Jia et al. 2004a). The Atf1-Pcr1 dimer binds to consensus sequences within the *mat* locus that exist at the *REIII* silencer and ∼1.4 k bp away from *REIII*, close to *cenH* (Fig. 1A) (Thon et al. 1999). Other histone-modifying enzymes such as histone deacetylases (HDACs) are also required for donor selection and heterochromatin establishment. These include the NAD^+^-dependent histone deacetylase Sir2 (Shankaranarayana et al. 2003) and the Snf/Hdac-containing repressor complex (SHREC), Clr1, Clr2 and Clr3 (Ekwall and Ruusala 1994; Thon and Klar 1992). Clr4, Clr3 and another HDAC, Clr6, interact with Atf1, therefore Atf1-Pcr1 has been suggested to recruit these histone modifying enzymes to the *mat* locus (Jia et al. 2004a; Kim et al. 2004; Yamada et al. 2005).

We recently conducted a genetic screen for factors that affect the directionality of mating-type switching. We identified the six genes (*set1, swd1, swd2, swd3, ash2* and *spf1*) encoding components of the H3K4 mono-, di- and tri-methyltransferase complex, Set1C/COMPASS. We also identified the *brl2* gene encoding a component of the H2BK119 monoubiquitin ligase HULC complex that is composed of four subunits, Brl1, Brl2, Rad6 and Shf1 (Maki et al. 2018; Thon et al. 2018). In *S. pombe*, no other enzyme catalyzes these reactions. Ubiquitylation of H2B (H2Bub at K119 in fission yeast, at K123 in budding yeast, and at K120 in human) is required for the tri-methylation of H3K4 by Set1/COMPASS family from yeast to human (Shilatifard 2012). Therefore, we hypothesized that H2Bub and H3K4me might work together to regulate MTS. Interestingly, the histone modifications by HULC and Set1C are generally observed at active genes, in euchromatin, and have been proposed to antagonize heterochromatin formation at the *mat* locus (Greenstein et al. 2020; Noma et al. 2001; Zofall and Grewal 2007). However, HULC and Set1C have also been reported to have a positive effect on gene silencing in heterochromatin (Chen et al. 2008; Kanoh et al. 2003; Mikheyeva et al. 2014). HULC was proposed to promote loading of heterochromatin factors in centromeric repeats by facilitating a wave of transcription that fuels the RNAi machinery during S-phase (Chen et al. 2008). In the case of Set1C, the deletion of the catalytic subunit Set1 causes a slight derepression of centromeric and telomeric reporter genes, and transcription of the *cenH* element in the *mat* region (Kanoh et al. 2003; Mikheyeva et al. 2014). In addition, Set1C also functions in the repression of the stress-response gene *ste11*(Materne et al. 2015; Materne et al. 2016) and *Tf2* retrotransposons (Mikheyeva et al. 2014). Interestingly, Set1 localizes with Atf1 binding sites at centromeres, where it contributes to the heterochromatin assembly with the Clr3 HDAC (Lorenz et al. 2014).

In this study, we performed genetic and molecular analyses of HULC, Set1C, and histone mutants to better understand the role of the HULC and Set1C complexes in mating-type switching. Our results support the view that, like Set1C (Maki et al. 2018), HULC affects donor choice by reducing the effectiveness of the *SRE3* enhancer in M cells and that H2B ubiquitylation by HULC and H3K4 methylation by Set1C operate in the same pathway for this function. Both modifying enzymes regulate Swi6 enrichment positively at the *SRE2* and *SRE3* elements. On the other hand, the two enzymes differentially affect heterochromatin formation and heterochromatic silencing at the *mat* locus. When combined with *dcr1*Δ, the *set1*Δ mutation caused a derepression of a reporter gene at *SRE3* while *shf1*Δ, a HULC mutant, did not. Thus, beyond increasing our understanding of donor selection through HULC and Set1C, our findings also shed light on gene silencing mechanisms by these complexes.

## Results

### The histone H2B ubiquitylation complex HULC is involved in MTS

Recently, we determined that a subunit of HULC, the *brl2*^*+*^ gene product, is required for efficient MTS by screening the gene deletion library, Bioneer version 5 (Maki et al. 2018). In an independent screen for MTS defects with version 2 of the same library, we identified another subunit of HULC, Shf1 (<u>S</u>mall <u>H</u>istone Ubiquitination <u>F</u>actor 1). Therefore, we systematically investigated the switching phenotypes of mutants in each of the four HULC subunits, the *shf1*Δ, *rad6*Δ, *brl1*Δ and *brl2*Δ mutants. MTS defects can be detected by measuring mating-type ratios in saturated liquid cell cultures, under conditions where the cells do not conjugate, because efficient switching results in an equal proportion of each cell type. Mutants in a few factors such as Swi2 mutants display variable mating-type ratios in independent cultures while other mutations uniformly bias cell populations towards P or M (Jakociunas et al. 2013; Maki et al. 2018). Thus, for each HULC mutant, four independent clones were constructed and analyzed to evaluate clonal variation by multiplex PCR. The wild-type *h*^*90*^ strain, PG4045, had nearly equal proportions of P and M cells, as expected, but all HULC mutants displayed a biased mating-type ratio which was around 33% P cells in all cultures examined (Fig. 2A and Supplementary Fig. S1A). The deletion mutants were also examined by iodine staining of plate-grown colonies, a classical assay for MTS efficiency based on the conjugation and subsequent sporulation of diploids. Colonies of the control PG4045 strain grown on MSA plate stained darkly with iodine, indicative of efficient MTS. The HULC mutants were less stained (Fig. 2B and Supplementary Fig. S1B) consistent with the observed cell-type bias. Double deletion of *shf1* and, respectively, *rad6, brl1* or *brl2* did not show any additive effect (Fig. 2C and Supplementary Fig. S1C). We concluded that HULC is involved in MTS.

**Fig 2.**
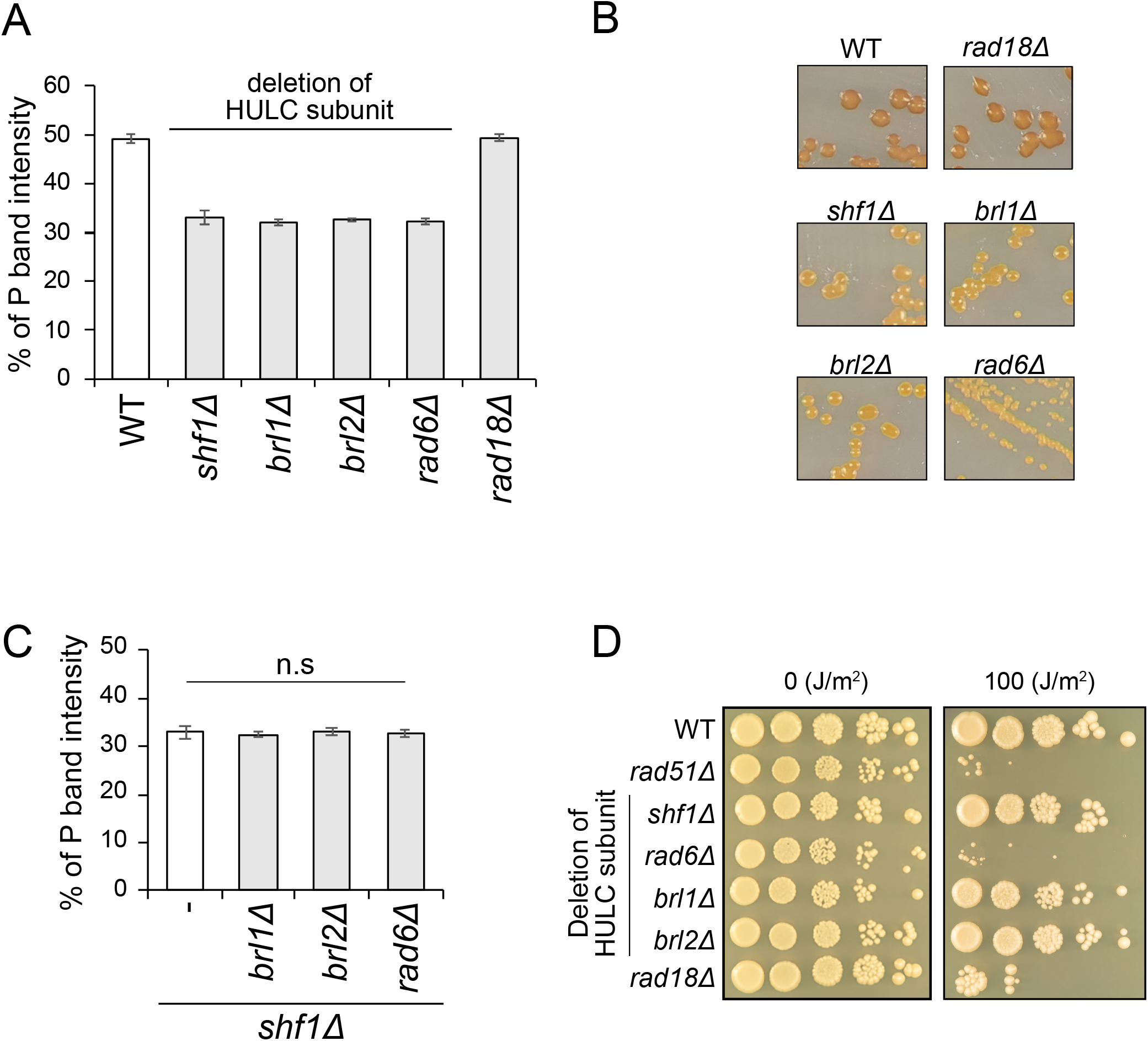
HULC is involved in MTS. (A) Multiplex PCR was used to measure the content of *mat1* using primers that bind specifically to either *mat1-P* or *mat1-M* together with a *mat1*-specific primer. The P and M band intensities were measured. The graph shows the mean value ± standard deviation (SD) for four independent colonies of a WT (*h*^*90*^) or single deletion mutants of each of the subunits of HULC and *rad18*. (B) Iodine staining of mutants lacking individual HULC subunits and a *rad18*Δ strain. WT (*h*^*90*^) colonies capable of forming spores stain darkly by exposure to iodine vapors due to the high content of starch within the spore wall. Colonies deficient in MTS stain lightly. (C) Quantification of *mat1* content estimated by multiplex PCR. Mean values ± SD of the % of P band intensity are displayed for four independent colonies of the single *shf1*Δ mutant (same samples as in A) and for double deletion mutants combining *shf1*Δ with deletions affecting other HULC subunits. Two-tailed paired Student’s *t* test was used to compare the mean of each sample to the control (the single *shf1*Δ mutant), n.s, not statistically significant. (D) Spot test using single deletion mutants of the HULC subunits and the *rad18*Δ mutant on a rich medium (YES) where a plate was subjected to UV radiation (100 J/m^2^). Five-fold dilution series are shown.

### The Rad18/Rad6 pathway does not influence MTS

In addition to its interaction with the E3 ubiquitin ligase Brl, the E2 Rad6 enzyme also operates with the E3 Rad18 in a conserved DNA damage tolerance pathway by mono-ubiquitylating PCNA (Hedglin and Benkovic 2015). We tested whether the Rad6/Rad18 pathway is required for MTS. As expected, both the *rad6* and *rad18* deletion mutant strains were sensitive to UV exposure, similar to a *rad51* deletion mutant strain (Fig. 2D). On the other hand, HULC mutants, respectively *shf1*Δ, *brl1*Δ and *brl2*Δ, did not show UV damage sensitivity (Fig. 2D). Conversely, multiplex PCR and iodine staining assays did not detect any switching defect in the *rad18*Δ mutant, unlike for the *shf1*Δ, *brl1*Δ and *brl2*Δ mutants (Fig. 2A and B, Supplementary Fig. S1A and S1B). These results indicate that Rad6 functions in different complexes for MTS and DNA damage tolerance.

### HULC regulates donor selection at *SRE3*

Depending on which step is affected, mutants deficient in MTS can be separated into three classes, Class Ia, Class Ib and Class II, by Southern blots (Egel et al. 1984). Class Ia mutants lack an imprint at *mat1*; the imprint is required for MTS as it is converted to a single-ended double-strand break just past the *H1* homology box during DNA replication, allowing invasion of *mat2-P* or *mat3-M* by the *H1* sequence (Fig. 1) (Klar et al. 2014; Thon et al. 2018). The imprint also creates a fragile site that causes breakage during DNA preparation and it is thus visible on Southern blots as a DNA double-strand break. Class Ib mutants have the imprint but fail to use it properly. Class II mutants are deficient at a later step, in the resolution of the gene conversion, which causes characteristic *mat2*-*mat3* cassette duplications inserted at *mat1* that are not found in Class I mutants. The *shf1* deletion was assigned to Class Ib in which a functional imprint is detected at the *mat1* cassette (Supplementary Fig. S2A) similar to the previously examined *brl2* deletion (Maki et al. 2018). The Class Ib group includes mutations that impair donor selection. These can be identified and further classified according to their effects in *h*^*09*^ cells that have swapped *mat2-M mat3-P* donor cassettes (Thon and Klar 1993). At least four groups of *h*^*09*^ Class Ib mutants can be distinguished: Group 1a displaying a strong bias towards P cells; Group 1b with a mild bias towards P cell; Group 2 with clonal variations; and Group 3 with a bias towards M cells (Maki et al. 2018). Deletions removing HULC subunits each caused a mild bias towards P cells in the *h*^*09*^ background (∼48 % P cells instead of ∼19 % in the presence of functional HULC), placing them in Group 1b (Fig. 3A and Supplementary Fig. S2B).

**Fig 3.**
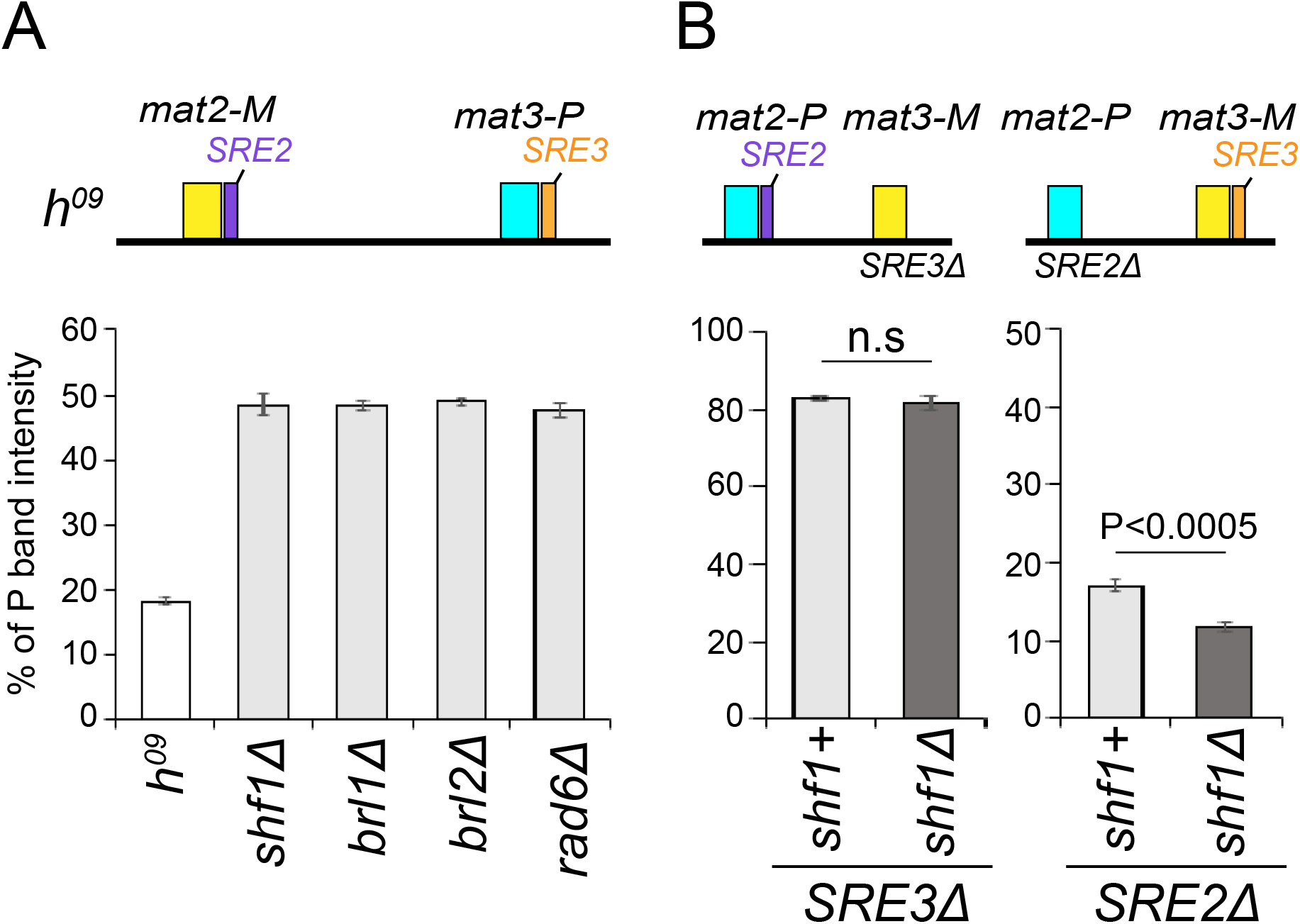
Analysis of donor preference in HULC mutants. (A) Quantification of *mat1* content in the *h*^*09*^ strain, of which the donor loci are swapped from *mat2-P mat3-M* to *mat2-M mat3-P*, by multiplex PCR. Mean values ± SD of the % of P band intensity of four independent colonies for single deletion mutants in each subunit of HULC in the *h*^*09*^ background. (B) Quantification of *mat1* content of mutants of *SRE* elements, *SRE3*Δ, *SRE2*Δ by multiplex PCR. Mean values ± SD of % of P band intensity of four or three independent colonies of strains with the mutated *SRE* element mutants, *SRE3*Δ and *SRE2*Δ. Two-tailed paired Student’s *t* test was used to compare the mean of each sample to control, n.s, not significant.

The *SRE* elements are necessary for donor selection (Jakociunas et al. 2013; Jia et al. 2004b). The changes in cell-type ratios caused by HULC mutations in *h*^*90*^(Fig. 2A; increased proportion of M cells) and *h*^*09*^ populations (Fig. 3A and Supplementary Fig. S2B; increased proportion of P cells) suggested in both cases an increased use of *SRE3* over *SRE2*. We further investigated the importance of *SRE2* and *SRE3* in the presence or absence of the *shf1* gene using *SRE* deletion mutants (Fig 3B). In an *SRE3*Δ strain, ∼84% of the population were P cells (i.e. use of *SRE2* is favored). This ratio was unaffected in *shf1*Δ *SRE3*Δ cells (∼83%) (Fig. 3B and Supplementary Fig. S2C). In an *SRE2*Δ strain, the*shf1* deletion slightly affected the mating-type ratio towards M cells (∼ 12% P cells in *shf1*Δ instead of ∼19 % in *shf1*^*+*^ cells) (Fig. 3B and Supplementary Fig. S2C). The slight shift in *shf1*Δ *SRE2*Δ cells does not seem very decisive, but it is statistically significant. These results suggest that, in M cells, HULC normally prevents the use of *SRE3*, rather than facilitating the use of *SRE2*.

### Histone residues modified by HULC and Set1C are required for MTS

We noticed that the genetic analyses in Fig. 3 showed similar trends for the MTS defects detected in HULC mutants and the defects previously reported for *set1* deletion mutants (Maki et al. 2018). The ubiquitylation of H2BK119 by HULC is essential for H3K4me3 by Set1C in *S. pombe* (Roguev et al. 2003), therefore we performed an epistasis analysis to test the relationship between HULC and Set1C in MTS. The switching defect of the *h*^*90*^ *shf1*Δ *set1*Δ double mutant was quite similar to the switching defect of the *h*^*90*^ *set1*Δ strain (∼42% P cells in *shf1*Δ *set1*Δ mutant *versus* ∼43% P cells in *set1*Δ mutant in the multiplex PCR assay), with *set1*Δ slightly suppressing the cell-type bias of the *shf1*Δ strain (∼33% P cells in *shf1*Δ mutant) (Fig. 4A and Supplementary Fig. S3A). This placed the two mutations in the same epistasis group, suggesting coordinated action of HULC and Set1C.

**Fig 4.**
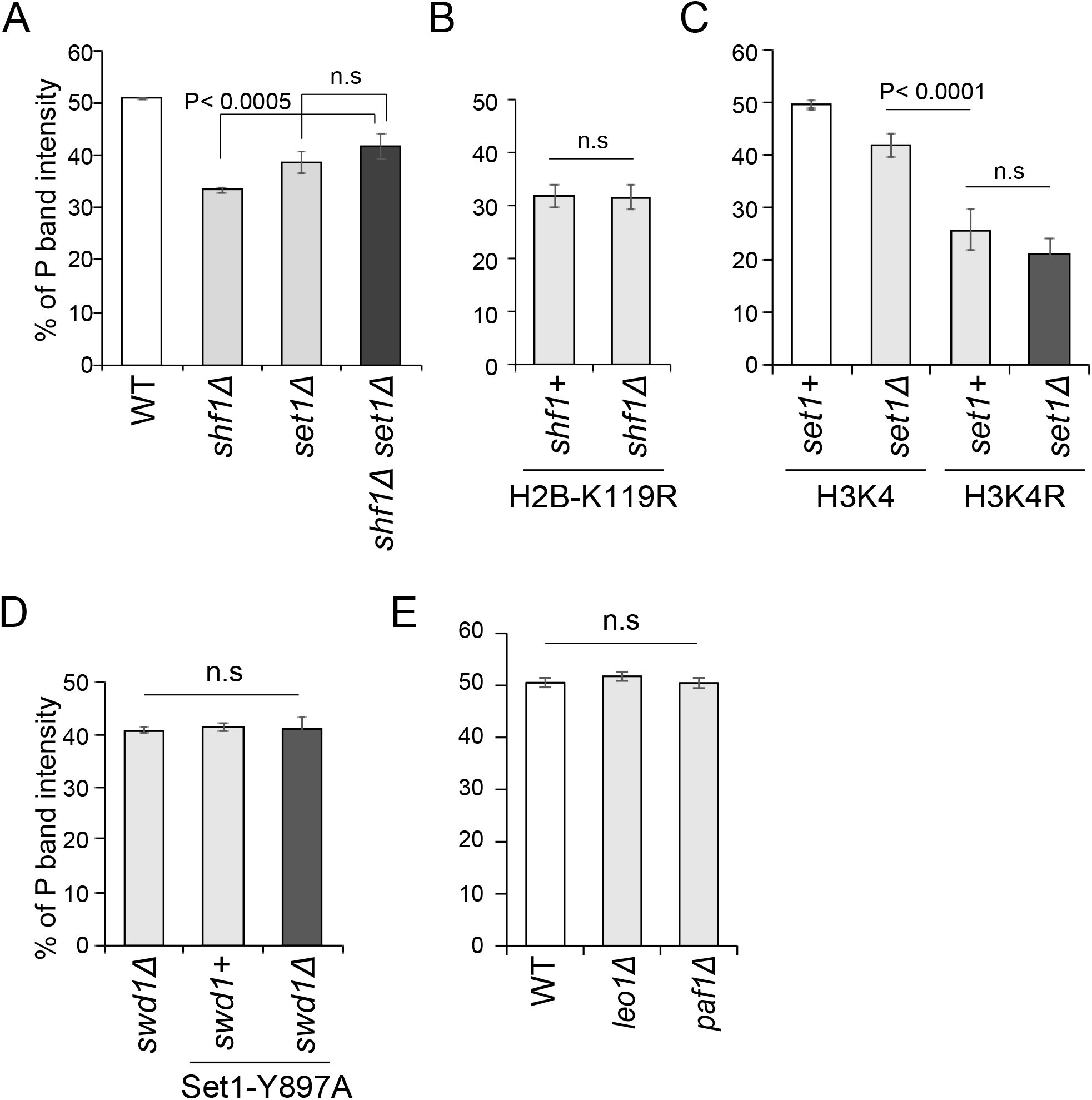
Analysis of histone modifications by HULC and Set1C in MTS. (A-E) Quantification of *mat1* content by multiplex PCR. Mean value ± SD of the % of P band intensity of four or six independent colonies, showing an epistasis analysis with *set1*Δ and *shf1*Δ (A), H2B-K119R mutation in an *shf1*^*+*^ or *shf1*Δ background (B), H3K4R mutation in a *set1*^*+*^ or *set1*Δ background (C), *swd1*Δ, the catalytically dead Set1 (Set1-Y897A) single mutant and *swd1*Δ Set1-Y897A double mutant (D) and deletion mutants of Paf1C subunits, *leo1*Δ and *paf1*Δ (E). Two-tailed paired Student’s *t* test was used to compare the mean of each sample to the WT control. “n.s”, not significant.

Next, we investigated the importance of the histone H2BK119 and H3K4 residues that are modified by HULC and Set1C, respectively, in MTS. The proportion of P cells in the H2BK119R mutant (∼34% P cells) was not only similar to the *shf1*Δ deletion mutant, but also to the double mutant *shf1*Δ H2BK119R (Fig. 4B and Supplementary Fig. S3B). On the other hand, the proportion of P cells in the H3K4R mutant was ∼27% P cells, less than in the *set1*Δ deletion mutant (∼44% P cells), but similar to the H3K4R *set1*Δ double mutant (∼30% P cells) (Fig. 4B and Supplementary Fig. S3C). We also investigated the requirement for the catalytic activity of Set1 in MTS. The SET domain has a highly conserved tyrosine residue which is suggested to be a catalytic residue from the crystal structure (Supplementary Fig. S3D) (Trievel et al. 2002), and that replacement of tyrosine Y1054 with alanine in *S. cerevisiae* Set1 causes loss of H3K4 methylation (Williamson et al. 2013). We created a Set1-Y897A mutant in *S. pombe*, corresponding to Set1-Y1054A in *S. cerevisia*e (Supplementary Fig. S3D). The mating-type ratios in the *set1-Y897A* mutant analyzed by multiplex PCR were similar to the ratios in the *set1*Δ mutant (Fig. 4D and Supplementary Fig. S3E). We verified that the Set1-Y897A protein tagged with 9×V5 epitope (9×V5-Set1-Y897A) was present in the cells (Supplementary Fig. S3F). The *swd1*^+^ gene encodes a subunit of the Set1C complex (Roguev et al. 2003). An *swd1*Δ single mutant and *swd1*Δ Set1-Y897A double mutant also showed similar ratios to the *set1*Δ mutant (Fig. 4D and Supplementary Fig. S3E). These observations point to both H2BK119 ubiquitylation by HULC and H3K4 methylation by Set1C playing an important role in the directionality of MTS.

Several lines of evidence have indicated that the RNA polymerase II-associated factor 1 complex Paf1C (Krogan et al. 2003; Ng et al. 2003; Wood et al. 2003), functionally conserved from yeast to mammals (Kim et al. 2009; Mbogning et al. 2013), can recruit HULC and Set1C (Sadeghi et al. 2015) and in fission yeast Paf1C prevents heterochromatin propagation across the *IR-L* boundary of the *mat* locus, among other effects (Sadeghi et al. 2015). In order to test the requirement for Paf1C in MTS, we constructed strains lacking the Paf1C components Leo1 and Paf1, respectively. The *leo1*Δ and *paf1*Δ mutants did not show a switching defect in the multiplex PCR assay (Fig. 4E and Supplementary Fig. S3G). Thus, the effects of HULC and Set1C in MTS occur independently of Paf1C.

### Set1 and Shf1 are involved in the mating-type-specific localization of Swi6 at *SRE2* and *SRE3*, but they have different effects on histone H3K9 methylation

The results presented so far allow updating a previous model (Maki et al. 2018) by now proposing that both HULC and Set1C reduce use of the *SRE3* recombination enhancer in M cells by modifying histone H2BK119 and H3K4. Given that differential Swi6 enrichment in the mating-type regions of M and P cells is a determinant of donor choice (Jakociunas et al. 2013; Jia et al. 2004b; Thon and Klar 1993), we next investigated the effects of HULC and Set1C on Swi6 occupancy at *SRE2, SRE3* and the K region located between *SRE2* and *cenH* (Fig. 5A). To this end, we performed ChIP-qPCR with Flag-tag antibody for 3×Flag-Swi6 using heterothallic strains with a fixed mating type, P (*mat1-PΔ17)* or M (*mat1-Msmt-0*). In a wild-type background, Swi6 showed an approximately three-fold higher enrichment at both *SRE2* and *SRE3* in M cells compared to P cells (Fig. 5B). No significant difference between P and M cells was observed within the K region, specifically at a location between *mat2-P* and *cenH* analyzed in this study (Fig. 5B). In both the *shf1*Δ and *set1*Δ backgrounds, the high, M-specific, Swi6 occupancy at *SRE2* and *SRE3* was decreased (Fig. 5B). It thus appears likely that Shf1 and Set1, and by extension HULC and Set1C, control donor choice at least in part by ensuring high Swi6 occupancy at recombination enhancers in M cells. We note, however, that the double *shf1*Δ *swi6*Δ deletion mutant showed a statistically significant slightly lower (in *h*^*90*^ cells) or higher (in *h*^*09*^ cells that have swapped *mat2-M mat3-P* donor cassettes) proportion of P cells compared with the single *swi6*Δ mutant (Supplementary Fig. S4), suggesting some effects of HULC in MTS are not through Swi6.

**Fig 5.**
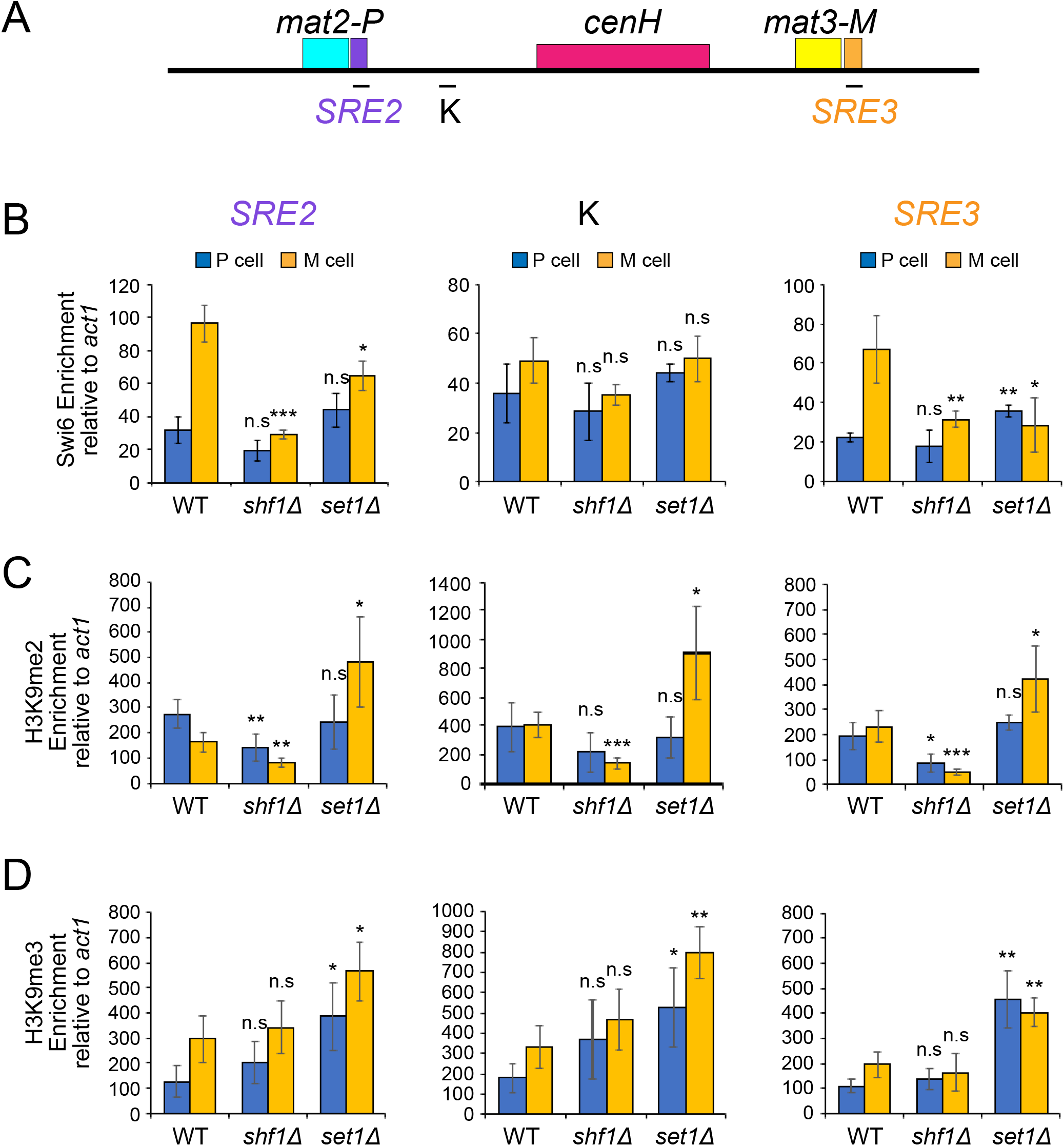
Shf1 and Set1 affect the enrichments of Swi6, H3K9me2 and -me3 in the *mat* silenced region. (A) Primer annealing sites used for qPCR: *SRE2*, K located between *SRE2* and *cenH*, and *SRE3*. (B-D) ChIP-qPCR analysis of enrichment levels relative to *act1* in P and M cells in WT, *shf1*Δ or *set1*Δ backgrounds for Swi6 (B), H3K9me2 (C) and H3K9me3 (D). Error bars indicate SD (n=3 or 4). Two-tailed paired Student’s *t* test was used to compare the mean obtained for each gene deletion to WT of the same mating-type, *P < 0.05; **P < 0.01; ***P < 0.001; n.s, not significant.

Swi6 recognizes di- and tri- methylation of histone H3K9 (Jih et al. 2017; Nakayama et al. 2001). We thus examined the di- and tri- methylation levels of H3K9 in *shf1*Δ and *set1*Δ strains using the same chromatin fixed samples as in Fig. 5B and antibodies specific for di- and tri- methylation of H3K9 (H3K9me2 and H3K9me3, respectively) (Fig. 5C and D). Comparing P and M cells, we observed that H3K9me2 enrichments in wild-type strains were comparable at the three locations examined and did not vary with cell type, with the possible exception of *SRE2* for which H3K9me2 was slightly lower in M cells (Fig. 5C). In *shf1*Δ strains, H3K9me2 was significantly reduced in both cell types with the similar tendency of slightly lower enrichments in M cells (Fig. 5C). The opposite trend was seen for *set1*Δ cells where H3K9me2 was unchanged in P cells but was increased in the case of M cells (Fig. 5C).

For H3K9me3, M cells showed somewhat higher enrichments than P cells in the wild-type background (Fig. 5D). This difference between P and M cells was attenuated in the *shf1*Δ background, but globally the levels of H3K9me3 enrichments remained unchanged in *shf1*Δ cells (Fig. 5D). As observed for H3K9me2 (Fig. 5C), H3K9me3 levels were increased by *set1*Δ, at all locations and in both cell types (Fig. 5D). These ChIP-qPCR analyses suggest that the enrichment levels of H3K9me2 and H3K9me3 are regulated by Set1, however these regulations work in different manners and correlations between H3K9 methylation and Swi6 enrichment levels were not systematically observed.

### Shf1 and Set1 have differential effects on silencing and MTS

Two main pathways of heterochromatin formation have been well documented at the *mat* locus, one is RNAi nucleating heterochromatin at the *cenH* region (Allshire and Ekwall 2015; Hall et al. 2002; Volpe et al. 2002), and the other is an RNAi-independent pathway involving the Atf1-Pcr1 complex (Greenstein et al. 2018; Jia et al. 2004a; Kim et al. 2004; Wang and Moazed 2017). Both H2Bub and H3K4me are believed to control transcription by regulating RNA polymerase II activity in *S. pombe* (Mikheyeva et al. 2014; Tanny et al. 2007; Zofall and Grewal 2007). This suggests that HULC and Set1C might participate in silencing the *mat* locus by facilitating RNAi which uses transcription products from *cenH* to nucleate heterochromatin. Alternatively, the fact that H2Bub and H3K4me mediate gene silencing at the *ste11* gene locus (Materne et al. 2015; Materne et al. 2016), together with the fact that Set1 localizes to Atf1 binding sites at centromeres and at the *ste11* gene locus (Lorenz et al. 2014), suggests that HULC and Set1C might co-operate with Atf1-Pcr1 also in the mating-type region. To investigate whether Shf1 or Set1 participates in the RNAi or Atf1-Pcr1 pathway of heterochromatin formation, we created double mutants, respectively *shf1*Δ *dcr1*Δ, *shf1*Δ *pcr1*Δ, *set1*Δ *dcr1*Δ and *set1*Δ *pcr1*Δ in a strain with a *ura4*^*+*^ reporter gene inserted in the *SRE3* region (PG1899 with *(EcoRV)*::*ura4*^*+*^) (Fig. 6A). As previously reported (Jia et al. 2004a), *dcr1*Δ and *pcr1*Δ single deletion mutants with the *(EcoRV)::ura4*^*+*^ reporter were resistant to 5-FOA, which is toxic to cells that express *ura4*^*+*^, while the *dcr1*Δ *pcr1*Δ double deletion caused sensitivity, at the same level as the *swi6*Δ strain (Fig. 6B). The *shf1*Δ single mutant was resistant to 5-FOA, and the *shf1*Δ *dcr1*Δ and *shf1*Δ *pcr1*Δ double mutants remained resistant. In contrast, the *set1*Δ and *set1*Δ *pcr1*Δ strains were resistant to 5-FOA, but the *set1*Δ *dcr1*Δ strain was clearly sensitive, its growth on 5-FOA was as severely affected as growth of the *dcr1*Δ *pcr1*Δ strain (Fig. 6B). This places *set1*Δ, but not *shf1*Δ, in the same epistasis group as *pcr1*Δ for the effects of these mutations on silencing in the mating-type region.

**Fig 6.**
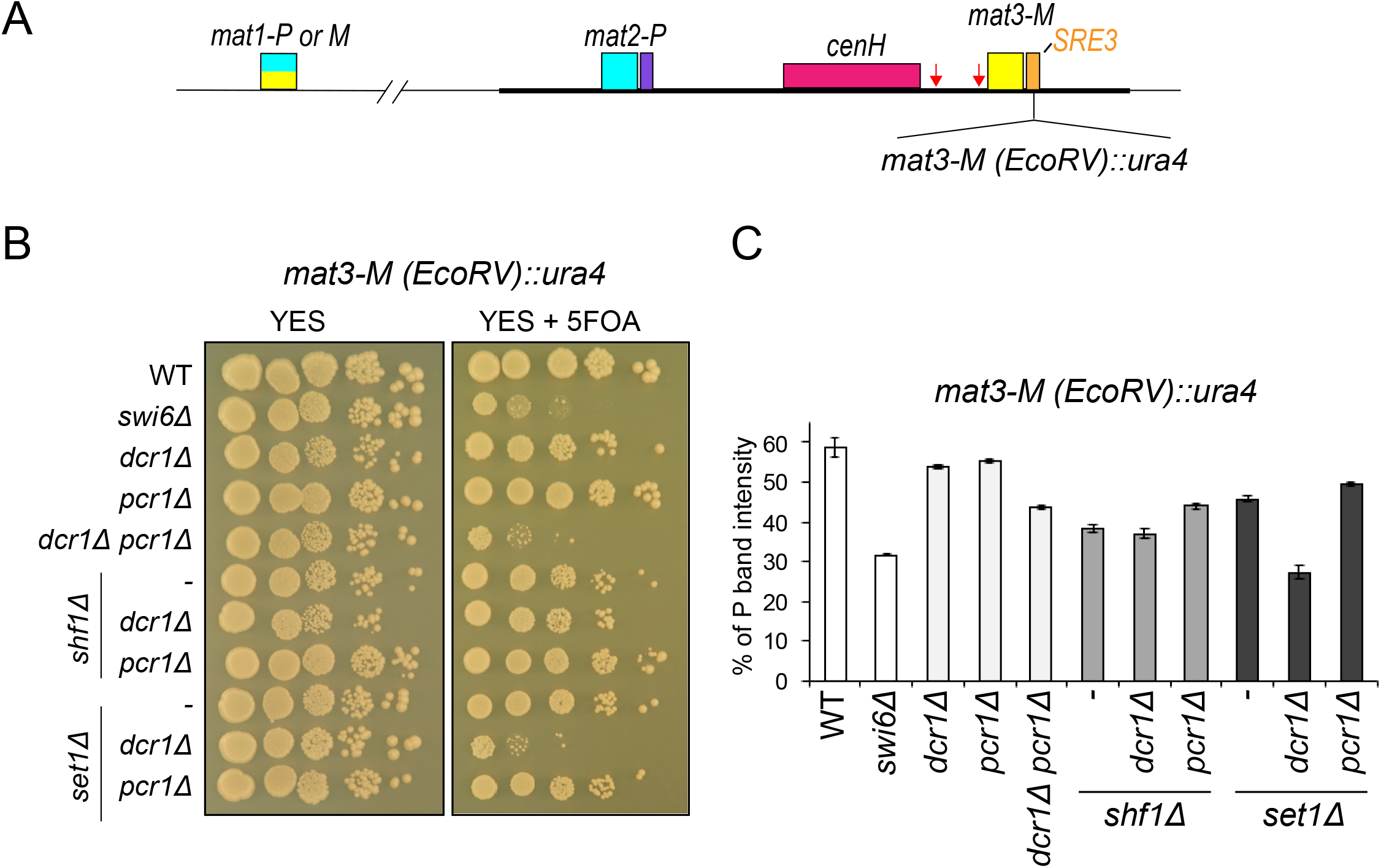
Set1 works in different pathway as Dcr1 for gene silencing and MTS. (A) Schematic view of the mating-type region of a *h*^*90*^ strain containing a *ura4* reporter gene (PG1899). Red arrows indicate binding sites for Atf1-Pcr1 dimers (B) Silencing in PG1899 was assayed by plating ten-fold dilution series of cells with the indicated gene deletions on YES medium or medium containing 5-FOA. (C) Quantification of *mat1* content by multiplex PCR, showing the % of P band intensity for the strains analyzed in the silencing assay in (B). Error bars indicate SD (n=3).

We examined the switching phenotype of the strains shown in Fig. 6B by multiplex PCR (Fig. 6C and Supplementary Fig. S5). The PG1899 strain carrying (*EcoRV)*::*ura4*^*+*^ at *SRE3* was slightly biased toward P cells (∼59% P cells) in comparison with the normal *h*^*90*^ configuration (∼50% P cells). This switching phenotype probably comes from the insertion of the *ura4*^*+*^ gene slightly affecting *SRE3* due to its proximity (see Fig. 6A). However, populations of a *swi6*Δ derivative of PG1899 showed mating-type biases similar to the *h*^*90*^ *swi6*Δ mutant, hence we analyzed all mutant strains derived from PG1899 for MTS. The single *dcr1*Δ and *pcr1*Δ mutants showed mating-type ratios similar to PG1899. The double *dcr1*Δ *pcr1*Δ mutant was biased towards M cells (∼45% P cells). The double *shf1*Δ *dcr1*Δ and *shf1*Δ *pcr1*Δ mutants had cell-type ratios similar to the *shf1*Δ single deletion mutant. In the case of *set1*Δ, the *set1*Δ *pcr1*Δ deletion strain had a cell-type ratio similar to the *set1*Δ strain, but the *set1*Δ *dcr1*Δ double deletion strain showed a larger bias toward M cells than the *set1*Δ strain, the largest bias of all strains examined. Thus, both gene silencing and multiplex PCR assays converge to show that Set1 functions in parallel to Dcr1 at the silent *mat* locus.

## Discussion

Mating-type switching is governed by chromatin conformation. Thus, the H3K9 methyltransferase CLRC and several HDACs have well documented roles in the formation of heterochromatin required for the directionality of switching. Our genetic screens identified euchromatic factors, the H2B ubiquitin ligase HULC and the H3K4 methyltransferase Set1C, as additional MTS factors (Fig. 2) (Maki et al. 2018). H3K4me3 has been put forward as a regulator of DSB formation promoting meiotic recombination in some eukaryotes (Serrano-Quilez et al. 2020; Tock and Henderson 2018). In the MTS of *S. pombe*, the lysine specific demethylases, Lsd1/Lsd2, are recruited to the *mat1* locus and involved in imprint formation (Holmes et al. 2012; Raimondi et al. 2018). On the other hand, we found that mutants in HULC or Set1C do not show an imprinting defect (Supplementary Fig. S2) (See also Maki et al., 2018). This places the action of HULC and Set1C at the donor choice step. It indicates that euchromatic histone modifications do not only function in DSB events, but also in the selection of donor templates by regulation of chromatin conformation. A potentially important feature is that these complexes engage in a cross-talk where H2B ubiquitylation upregulates H3K4 methylation by Set1C (Tanny et al. 2007). Here, we obtained evidence that HULC and Set1C function in a common pathway for MTS to inhibit the preferential use of the *SRE3* enhancer in M cells, plausibly through this cross-talk. Notably, the effects of the two complexes determined by mutational analyses were not identical; there are distinct contributions to gene silencing detected for Set1 and Shf1 that likely reflect some divergent functions in heterochromatin formation.

Strong biases towards the M cell-type were observed in both HULC and Set1C mutants denoting increased use of the *SRE3* enhancer in these mutants (Fig. 2 and 3) (Maki et al. 2018). An epistasis analysis assigned the two complexes to the same pathway (Fig. 4), even though a slightly more pronounced bias towards M cells was observed in *shf1*Δ cells than in *set1*Δ or *shf1*Δ *set1*Δ cells (Fig. 4A). It has been reported that Set1 is still expressed in cells that lack HULC and low levels of ubiquitylation of H2BK119 *in vivo* (Mikheyeva et al. 2014; Tanny et al. 2007; Zofall and Grewal 2007) and H3K4 di-methylation are still detected in an H2BK119R strain (Tanny et al. 2007). In *S. cerevisiae*, H3K4 mono-methylation is still detected in *rad6* deletion cells (Schneider et al. 2005). Therefore, we speculate that the residual mono- and/or di- methylation of H3K4 in cells lacking H2Bub might increase the bias towards M cells, either directly through effects on recombination or indirectly. Our study also revealed that while the H2BK119R, *shf1*Δ and combined H2BK119R *shf1*Δ mutations had the same effect on MTS (Fig. 4B), the switching defects in the H3K4R and H3K4R *set1*Δ mutants were much stronger than in the single *set1*Δ strain (Fig. 4C). Transient acetylation of H3K4 has been proposed to facilitate the association of Swi6 with histone H3K9me2 through a chromodomain switch where Swi6 replaces H3K9me2-bound Chp1 or Clr4 at centromeres (Xhemalce and Kouzarides 2010). The same mechanism facilitating Swi6 association at *SRE* elements at the *mat* locus might account for the pronounced effects of H3K4R on MTS where Swi6 association is paramount.

The differential association of Swi6 with the mating-type region is a distinguishing factor between P and M cells: Swi6 is present at a low level over the region in P cells, coinciding with Swi2 specifically at *SRE3*, but at a high level in M cells, coinciding with Swi2 at both *SRE2* and *SRE3* (Jia et al. 2004b). These associations are believed to favor the use of *SRE2* over *SRE3* in M cells (Jakociunas et al. 2013; Yu et al. 2012). We found here that Shf1 (and to a lesser extent Set1) is required for the high Swi6 occupancy at *SRE2* and *SRE3* in M cells. In the *shf1*Δ and *set1*Δ mutants, Swi6 occupancy remained abnormally low at both enhancers in M cells, similar to what is normally seen in P cells (Fig. 5). This profile could account for *SRE3* being preferred over *SRE2* in both cell types in the mutants, where all cells would in essence behave as P cells. The association of Swi6 with the enhancers was most strongly reduced in the *shf1*Δ mutant (Fig. 5), where donor choice was also most impaired (Fig. 4). These effects would place HULC and Set1C upstream of the enhanced Swi6 association with recombination enhancers, *SRE2* and *SRE3*, in M cells (Fig. 7), without excluding other points of action. We propose that the resultant chromatin organization promotes *mat2-P* donor choice at *SRE*2 (Fig. 7).

**Fig 7.**
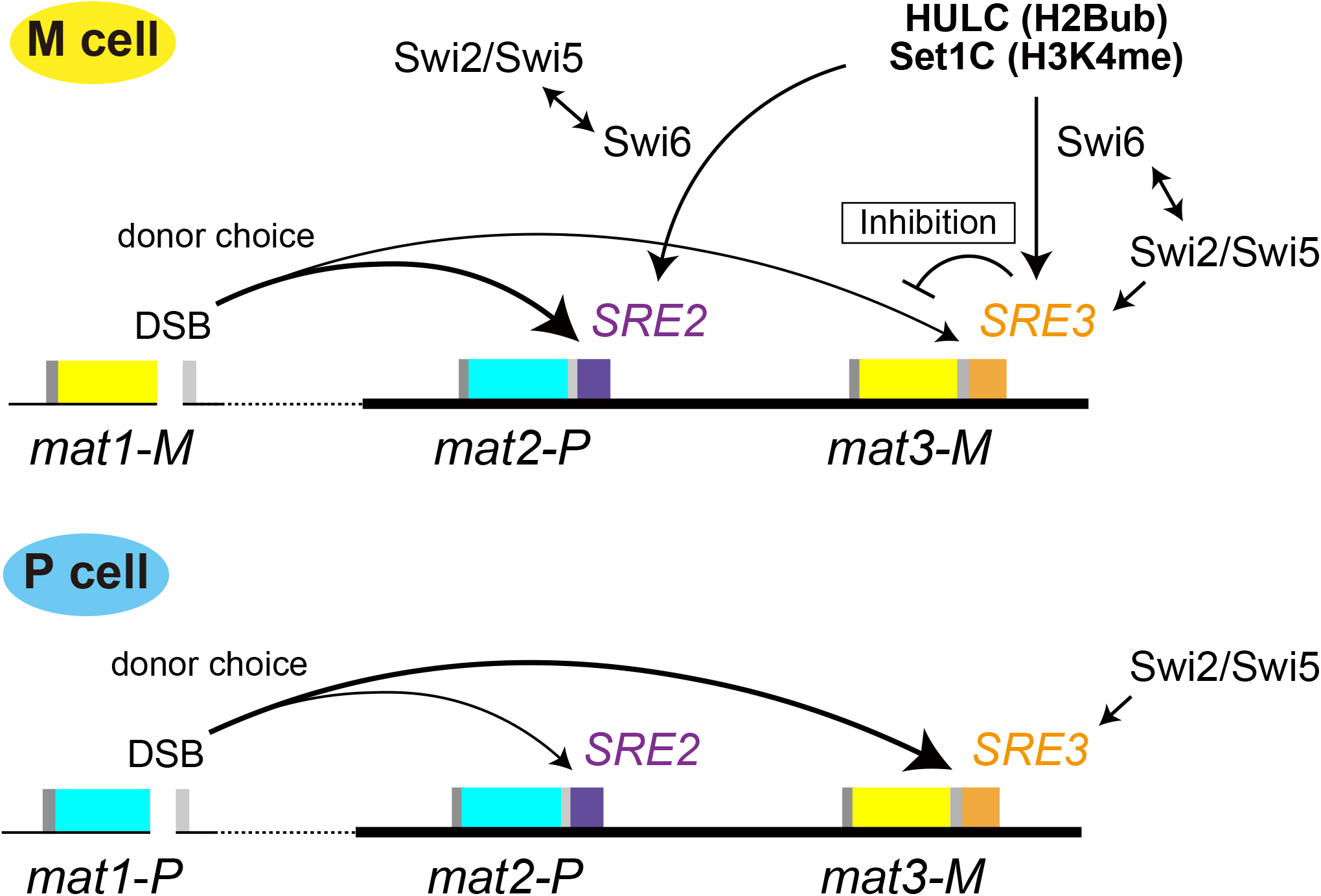
Proposed working model for regulation of MTS involving HULC and Set1C. In M cells, HULC and Set1C inhibit the *mat3-M* donor choice at *SRE3*. Swi6 enrichments are also regulated by HULC and Set1C at *SRE2* and *SRE3*. The Swi2-Swi5 complex associates with *SRE2* and *SRE3* in a Swi6-dependent or -independent manner. The resultant heterochromatin organization may promote *mat2-P* donor choice at *SRE2* by an unknown mechanism. In P cells, the Swi2-Swi5 complex localizes at *SRE3* specifically and promotes *mat3-M* donor choice, as previously suggested by Maki et al (2018).

Our experiments are consistent with and support an active repression by HULC and Set1C at *SRE3*, relevant to the central question of how *SRE2* outcompetes *SRE3* in M cells when Swi2 is present at both enhancers. *SRE2* might be inherently more efficient than *SRE3* under the conditions, or recombination might be actively repressed at *SRE3*. Here, *shf1*Δ and *set1*Δ mutants showed a bias towards M cells in *h*^*90*^ cells (Fig. 2 and 4) and towards P cells in *h*^*09*^ cells (Fig. 3A), consistent with the two factors repressing use of *SRE3*. Moreover, the ∼80% bias towards P in *SRE3*Δ cells remained unchanged in the *shf1*Δ mutant, indicating *SRE2* is functional in the absence of Shf1. In contrast, in the *SRE2*Δ strain, *SRE3* was increasingly used in *shf1*Δ cells, indicating Shf1 inhibits use of *SRE3* (Fig. 3B). Thus, our genetic analyses suggest that HULC inhibits selection of *SRE3* - and thereby *mat3-M* donor selection - in *h*^*90*^ cells, similar to Set1C (Fig. 7) (Maki et al. 2018).

How might an inhibition of recombination by HULC/Set1C take place at *SRE3*? A relevant effect of HULC/Set1C could be by controlling nucleosome occupancy or positioning. In *S. cerevisiae*, nucleosome occupancy is decreased genome-wide in H2BK123R, *rad6*Δ, and *lge1*Δ mutants (Batta et al. 2011; Chandrasekharan et al. 2009). In *S. pombe*, H2Bub decreases chromatin remodeling by RSC at the *ste11* promoter to repress transcription (Materne et al. 2015; Materne et al. 2016). In this case, the effect of H2Bub is through H3K4me and histone deacetylation, supporting the idea that HULC, Set1C, and HDACs might work in concert to position nucleosomes at the *mat* locus as well. Nucleosome positioning at *SRE3* might mask the enhancer or prevent strand invasion. An intriguing alternative that our results do not exclude is that nucleosome positioning or modification by HULC/Set1C might facilitate recombination near *SRE2* in M cells. Positive effects on recombination repair and on recombination-dependent bypass of DNA lesions have been reported for the RNF20/RNF40 mammalian homolog of Bre1 (Moyal et al. 2011; Nakamura et al. 2011) and for the *S. cerevisiae* counterpart of HULC (Hung et al. 2017). Nucleosome depletion by associated remodelers, rather than nucleosome stabilization, often appears instrumental during repair, highlighting the context dependency of the effects of the modifying and remodeling complexes (Challa et al. 2021; Nakamura et al. 2011). It will be important in the future to understand how remodeling complexes might contribute to MTS, taking into account the fact that Swi6 also interacts with many remodelers (Motamedi et al. 2008).

Finally, we uncovered effects of HULC and Set1C on the heterochromatic structure of the mating-type region that bring light to how these complexes might affect Swi6 association and MTS. In the case of *shf1*Δ, we observed reduced H3K9me2 at the three locations tested (Fig. 5C). HULC has been suggested to associate with the RNAi machinery in centromeric regions when repetitive sequences are transcribed during S-phase (Chen et al. 2008), by analogy *cenH* in the mating-type region might be an entry point for HULC. In the case of *set1*Δ, we observed a synthetic silencing defect when combining the *set1*Δ and *dcr1*Δ mutations (Fig. 6B), showing that Set1 and Dcr1 participate in parallel pathways of heterochromatin formation. The effect is clearly relevant to MTS as the double mutant showed a strong bias towards M cells (Fig. 6C). Similarly, Dcr1 and the Clr3 HDAC operate in parallel to recruit Clr4 to the *mat* locus, Dcr1 through RNAi at *cenH* and Clr3 through the Atf1-Pcr1 binding sites (Yamada et al. 2005). Here, we placed *set1*Δ in the same epistasis group as *pcr1*Δ for the effects of these mutations on silencing (Fig. 6B), suggesting that the Atf1-Pcr1 binding sites at the *mat* locus constitute entry points not just for Clr3 but also for Set1C. It has been suggested that H3K9me2 is associated with RNAi machinery and the transition from H3K9me2 to -me3 is associated with an RNAi-independent mechanism in centromeric regions (Jih et al. 2017). Distinct effects of *shf1*Δ and *set1*Δ on modifications of H3K9 and silencing of the *ura4* reporter gene inserted at *SRE3* may due to the differential functions of HULC and Set1C for RNAi machinery in the *cenH* region and RNAi-independent machinery using Atf1/Pcr1 complex. Independently, a genome-wide study found that Set1 cooperates with Clr3 to repress transcription at other Atf1-Pcr1 binding sites (Lorenz et al. 2014) and an important effect of Clr3 in heterochromatin is to suppress histone turnover (Aygun et al. 2013; Greenstein et al. 2018; Wang and Moazed 2017). Thus, taken altogether, HULC and Set1C might be recruited in several ways to the *mat* locus where they would cooperate with Clr3 to regulate aspects of heterochromatin formation and nucleosome occupancy important for donor selection. To further analyze this mechanism, we will need to understand how the regulation would be exerted in a cell-type-specific manner, as well as spatially, whether the regulation has to occur locally at the enhancers, or whether global effects on nucleosome mobility that would differ in P and M cells might lead to the observed biases in enhancer use.

## Material and Methods

### Yeast strains, strains constructions and strain manipulations

*S. pombe* strains used in this study are described in Supplementary Table S1. Standard techniques were used to cultivate, sporulate, cross and genetically manipulate *S. pombe* (Ekwall and Thon 2017). Strains were generated by transformations or genetic crosses. The H3K4R mutant strain (TM504: *h*^*90*^ *hht1-H3K4R hht2-H3K4R leu1-32 his3-D1*) was generated from EM20 (*h*^*90*^ *leu1-32 his3-D1 ade6-M375*) by CRISPR/Cas9-mediated gene editing. EM20 was transformed by a gRNA expression plasmid (pEM59), a Cas9 expression plasmid and HR donor templates. To mutate the *hht1 and hht2* genes, the common target sequence, 5’-TCTACCGGTGGTAAGGCACC-3’, was inserted into the BbsI site in gRNA scaffold sequence in pEM59. To select cells in which Cas9 was active, the *ade6-M375* mutation was edited in the same transformation. The gRNA targeting *ade6-M375*, 5’ CCTGCCAAACAAATTGATTG was also expressed from pEM59. The HR donor templates for mutagenesis, purchased from Integrated DNA Technologies, were amplified using primer sets, *hht1-H3K4R:* (5’-CTGCAGTACGCTTGCGTTTC-3’ and 5’-GGACGATAACGATGAGGCTTC-3’), *hht2-H3K4R* (5’-GGGAAGCCGAAATCGCAATC-3’ and 5’-CCAGGACGATAACGATGAGGCTTC-3’) and *ade6*^*+*^ (5’-GTGGTCAATTGGGCCGTAT-3’ and 5’-CGTGCACTTCTTAGACAGTTCA-3’) by PCR. Cells were grown on low-adenine plates and white colonies (*ade6*^*+*^) were selected. TM863 and TM896 were derived from TM504. TM501 was also generated by Cas9-mediated gene editing, with the gRNA target sequence (5’-TACTTATGATTACAAGTTTC-3’) and the HR donor template amplified by primer set (5’-GGGAAATATCGCGCGTTTC-3’ and 5’-CTAGTTTAAATAGCCACGACATGT-3’). All mutants selected for further analysis were confirmed by PCR and sequence analysis. The plasmid pEM59, allowing expression of an inserted gRNA sequence, was constructed by Emil Damgaard Jensen in the Thon lab. The sequence data is provided in Supplementary Fig. S6. The plasmid is available from Dr. Thon upon request. The sequences of the PCR-amplified HR donor templates are shown in Supplementary Fig. S7.

### Iodine staining

Estimates of the efficiency of MTS were obtained by iodine staining as described previously (Thon and Klar 1993). Briefly, cells were streaked on MSA plates, grown at 26°C for 3∼4 days. The plate on which the colonies were formed was turned upside down over a cylinder glass (dia. 90 mm) out with a few crystals of iodine on its bottom in a fume hood. The iodine vapors stained the colonies in about 10 min at room temperature.

### Multiplex PCR

Multiplex PCR was performed as previously described (Maki et al. 2018). *S. pombe* cells were propagated in 2 ml YE5S cultures at 30°C to saturation. Each culture (500 μl) was harvested in a 1.5 ml microcentrifuge tube, followed by DNA extraction with a Dr. GenTLE (from yeast) high recovery kit (Takara Bio). The genomic DNA concentration was measured using the Quantifluor® ONE dsDNA Dye System (Promega) and genomic DNA (5∼20 ng) was added PCR reaction mixture (total 20 μl). The primers used were FAM-MT1 (5’-AAATAGTGGGTTAGCCGTGAAAGG-3’) at 400 nM, MP1 (5’-ATCTATCAGGAGATTGGGCAGGTG-3’) at 200 nM and MM1(5’-GGGAACCCGCTGATAATTCTTGG-3’) at 200 nM. The 5’ end of FAM-MT1 and FAM-MT3 was labeled with 6-carboxyfluorescein (FAM). To reduce non-specific PCR products, 400 nM heat-stable RecA protein from *Thermus thermophiles* and 400 μM ATP were included in the PCR reaction buffer (10 mM Tris-HCl pH 8.3, 50 mM KCl, 2.5 mM MgCl2) (Shigemori et al. 2005). The amplification program was used: 2 min at 94°C - 27 x [30 sec at 94°C - 30 sec at 55°C - 1 min at 72°C] - 5 min at 72°C. PCR fragments corresponding to *mat1-P*and *mat1- M* alleles were resolved on 5% polyacrylamide gels. Fluorescence was detected and quantified using Typhoon FLA9500 (GE Healthcare) and ImageQuant (GE Healthcare).

### UV damage sensitivity

Serial dilutions of exponentially growing cell cultures were plated on complete medium (YES) and subjected to UV irradiation by exposure to a germicidal lamp (254 nm; 100 J/m^2^). UV intensities were measured with a UV Radiometer (TOPCON UVR-2). Plates were incubated at 30°C for 3∼4 days.

### Chromatin Immunoprecipitation (ChIP)

ChIP was performed as in Kimura et al. (2008), with a few modifications. EMM2 medium (50 mL) containing 0.1 g/l each of leucine, adenine, histidine, uracil and arginine was used for cell culture. The cultures were propagated to 1.0 × 10^7^ cells/ml at 30°C, and then shifted to 18°C for 2 h. 5.0 × 10^8^ cells were cross-linked with 1% formaldehyde for 15 min at 25°C and then incubated in 125 mM glycine for 5 min. Cross-linked cell lysates were solubilized by a multi-beads shocker (Yasui Kikai) at 4°C, 15 cycle of 1 min on and 1 min off, and sonicated using a Bioruptor UCD-200 (Diagenode) at 2 cycle of 10 min each with alternating pulses of 40 sec on and 30 sec off at high level. The sheared samples were centrifuged at 20,000 g for 10 min at 4°C. The supernatants were incubated with 30 μl Dynabeads Protein A (Thermo Fisher) preloaded with 1.2 μl anti-FLAG M2 antibody (Sigma-Aldrich) for 6 h at 4°C. The beads were washed sequentially with wash buffer 1 (50 mM Hepes-KOH [pH7.5], 1 mM EDTA, 0.1 % Sodium deoxycholate, 0.5 M NaCl), wash buffer 2 (10 mM Tris-HCl [pH8.0], 0.25 M LiCl, 1 mM EDTA, 0.5% NP40, 0.5% SDS) and TE (twice), and materials coprecipitated with the beads were eluted with elution buffer (50 mM Tris-HCl [pH 7.6], 10 mM EDTA and 1% SDS) for 20 min at 65°C. The eluates were incubated at 65°C overnight to reverse cross-links and were then treated with 10 μg/ml RNase A for 1 hr at 37°C, followed by 20 μg/ml proteinase K for 3 hr at 50°C. DNA was purified with a MonoFas DNA purification kit I (GL Sciences). Quantitative PCR was performed with SYBR Premix Ex Taq II (TaKaRa Bio) or TB Green Premix DimerEraser (Takara Bio) on a Mx3000P qPCR system (Agilent). Primer sequences are in Supplementary Table S2. For ChIP of H3K9me2 and -me3, 20 μg of H3K9me2 antibody or 20 μg of H3K9me3 antibody (Hayashi-Takanaka et al. 2011) were preloaded to 40 μl Dynabeads M-280 Sheep anti-Mouse IgG (Thermo Fisher).

## Acknowledgments

We are grateful to members of the Iwasaki Laboratory for discussion. We thank Jason Tanny for providing us with the *htb1* mutant strain and Jun-ichi Nakayama for providing us with the Flag-Swi6 strain. We thank the Biomaterials Analysis Division, Open Facility Center, Tokyo Institute of Technology for sequence analysis.

## Author contributions

A.E.C. and T.M. conducted experiments. T.M. and H.I. were responsible for conceptualization and project design. T.M., H.T., T.H., H.K., G.T. and H.I. supervised the study. H.K. provided materials. A.E.C., T.M., J.E.H, G.T and H.I. wrote the manuscript. T.M., J.E.H, G.T and H.I. were responsible for data analysis and funding acquisition.

