## supplementary info for "Euchromatin factors HULC and Set1C affect heterochromatin organization for mating-type switching in fission yeast *Schizosaccharomyces pombe*"

**By**

**Alfredo Esquivel-Chávez, Takahisa Maki, Hideo Tsubouchi, Testuya Handa, Hiroshi Kimura, James E. Haber, Genevieve Thon and Hiroshi Iwasaki**

Supplementary Materials including
Supplementary Tables 1 and 2
Supplementary Figures S1- S7
Supplementary Materials and Methods

### Supplementary **Table S1. Lest of strains used in this study**

| **Strain** | **Mating type region** | **Genotype** |
|---|---|---|
| PG4045 | $h^{90}$ | *(Blp1)::LEU2 leu1::*$ura4^{+}$*-[mfm3p-YFP]-[map2p-CFP] ura4-D18 ade6-M216* |
| PAE0207 | $h^{90}$ | *(Blp1)::LEU2 leu1::*$ura4^{+}$*-[mfm3p-YFP]-[map2p-CFP] ura4-D18 ade6-M216 shf1Δ::kamR* |
| PAE0302 | $h^{90}$ | *(Blp1)::LEU2 leu1::*$ura4^{+}$*-[mfm3p-YFP]-[map2p-CFP] ura4-D18 ade6-M216 brl1Δ::kamR* |
| PAE0450 | $h^{90}$ | *(Blp1)::LEU2 leu1::*$ura4^{+}$*-[mfm3p-YFP]-[map2p-CFP] ura4-D18 ade6-M216 brl2Δ::kamR* |
| PAE0905 | $h^{90}$ | *(Blp1)::LEU2 leu1::*$ura4^{+}$*-[mfm3p-YFP]-[map2p-CFP] ura4-D18 ade6-M216 rhp6Δ::kamR* |
| PAE0061 | $h^{90}$ | *(Blp1)::LEU2 leu1::*$ura4^{+}$*-[mfm3p-YFP]-[map2p-CFP] ura4-D18 ade6-M216 rhp6Δ::kamR shf1Δ::hphR* |
| PAE1101 | $h^{90}$ | *(Blp1)::LEU2 leu1::*$ura4^{+}$*-[mfm3p-YFP]-[map2p-CFP] ura4-D18 ade6-M216 brl1Δ::kamR shf1Δ::hphR* |
| PAE1102 | $h^{90}$ | *(Blp1)::LEU2 leu1::*$ura4^{+}$*-[mfm3p-YFP]-[map2p-CFP] ura4-D18 ade6-M216 brl2Δ:: kamR shf1Δ::hphR* |
| PAE0018 | $h^{90}$ | *(Blp1)::LEU2 leu1::*$ura4^{+}$*-[mfm3p-YFP]-[map2p-CFP] ura4-D18 ade6-M216 rad18Δ::kamR* |
| PG4048 | $h^{09}$ | *(Blp1)::LEU2 leu1::*$ura4^{+}$*-[mfm3p-YFP]-[map2p-CFP] ura4-D18 ade6-M216* |
| PAE0210 | $h^{09}$ | *(Blp1)::LEU2 leu1::*$ura4^{+}$*-[mfm3p-YFP]-[map2p-CFP] ura4-D18 ade6-M216 shf1Δ::kamR* |
| PAE0305 | $h^{09}$ | *(Blp1)::LEU2 leu1::*$ura4^{+}$*-[mfm3p-YFP]-[map2p-CFP] ura4-D18 ade6-M216 brl1Δ::kamR* |
| PAE0304 | $h^{09}$ | *(Blp1)::LEU2 leu1::*$ura4^{+}$*-[mfm3p-YFP]-[map2p-CFP] ura4-D18 ade6-M216 brl2Δ::kamR* |
| PAE0906 | $h^{09}$ | *(Blp1)::LEU2 leu1::*$ura4^{+}$*-[mfm3p-YFP]-[map2p-CFP] ura4-D18 ade6-M216 rhp6Δ::kamR* |
| TP75 | *mat3-M SRE3Δ* | *ura4-D18 leu1-32 ade6-M210* |
| PAE1258 | *mat3-M SRE3Δ* | *ura4-D18 leu1-32 ade6-M210 shf1Δ::kamR* |
| TP8 | *mat2-P SRE2Δ* | *ura4-D18 leu1-32 ade6-M210* |
| TM918 | *mat2-P SRE2Δ* | *ura4-D18 leu1-32 ade6-M210 shf1Δ::hphR* |
| TP126 | *mat3-M-SRE2* | *ura4-DS/E leu1-32 ade6-216* |
| PAE127 | *mat3-M-SRE2* | *ura4-DS/E leu1-32 ade6-216 shf1Δ::kamR* |
| TP303 | *mat2-P-SRE3* | *ura4-D18 leu1-32 ade6-210* |
| PAE304 | *mat2-P-SRE3* | *ura4-D18 leu1-32 ade6-210 shf1Δ::kamR* |
| TM213 | $h^{90}$ | *3xFlag-swi6, leu1-32, ura4-d18* |
| PAE1937 | $h^{90}$ | *3xFlag-swi6 leu1-32 ura4-D18 shf1Δ:: kamR* |
| PAE1421 | $h^{90}$ | *3xFlag-swi6 leu1-32 ura4-D18 set1Δ:: kamR* |
| TM865 | $h^{90}$ | *3xFlag-Swi6 leu1-32 set1D::kanR shf1D::hphR* |
| PAE4046 | $h^{90}$ | *(Blp1)::LEU2 leu1::*$ura4^{+}$*-[mfm3p-YFP]-[map2p-CFP]* |

| | | |
|---|---|---|
| | | *ura4-D18 ade6-M216 leo1Δ::kamR* |
| PAE4047 | $h^{90}$ | *(Blp1)::LEU2 leu1::*$ura4^{+}$*-[mfm3p-YFP]-[map2p-CFP] ura4-D18 ade6-M216 paf1Δ::kamR* |
| PAE048 | $h^{90}$ | *(Blp1)::LEU2 leu1::*$ura4^{+}$*-[mfm3p-YFP]-[map2p-CFP] ura4-D18 ade6-M216 htb1-K119R::kamR* |
| PAE0757 | $h^{90}$ | *(Blp1)::LEU2 leu1::*$ura4^{+}$*-[mfm3p-YFP]-[map2p-CFP] ura4-D18 ade6-M216 htb1-K119R:: kamR shf1Δ::hphR* |
| TM899 | $h^{90}$ | *hht3D::hphR leu1-32 his3-D1* |
| TM912 | $h^{90}$ | *hht3D::hphR (Blp1)::LEU2 leu1::*$ura4^{+}$*-[mfm3p-YFP]-[map2p-CFP] ura4-D18 ade6-M216 set1Δ::kamR* |
| TM863 | $h^{90}$ | *hht1- H3K4R, hht2-H3K4R hht3D::hphR leu1-32 his3-D1* |
| TM896 | $h^{90}$ | *hht1- H3K4R, hht2-H3K4R hht3Δ::hphR leu1-32 his3-D1 set1Δ::kanR* |
| TM501 | $h^{90}$ | *set1-Y897A leu1-32 his3-D1* |
| TM906 | $h^{90}$ | *set1-Y897A swd1D::kamR leu1-32 his3-D1* |
| TM777 | $h^{90}$ | *9xV5-set1-Y897A::hphR* |
| TM214 | *mat1-PΔ17::LEU2* | *3Flag-swi6 leu1-32 ura4-D18* |
| TM215 | *mat1-Msmt-0* | *3Flag-swi6 leu1-32 ura4-D18* |
| PAE1025 | *mat1-PΔ17::LEU2* | *3Flag-swi6 leu1-32 ura4-D18 shf1Δ::kamR* |
| PAE1023 | *mat1-Msmt-0* | *3Flag-swi6 leu1-32 ura4-D18 shf1Δ::kamR* |
| PAE1985 | *mat1-PΔ17::LEU2* | *3Flag-swi6 leu1-32 ura4-D18 set1Δ::kamR* |
| PAE1915 | *mat1-Msmt-0* | *3Flag-swi6 leu1-32 ura4-D18 set1Δ::kamR* |
| PAE006 | $h^{90}$ | *(Blp1)::LEU2 leu1::*$ura4^{+}$*-[mfm3p-YFP]-[map2p-CFP] ura4-D18 ade6-M216 swi6Δ::kamR* |
| PAE092 | $h^{90}$ | *(Blp1)::LEU2 leu1::*$ura4^{+}$*-[mfm3p-YFP]-[map2p-CFP] ura4-D18 ade6-M216 swi6Δ::kamR shf1Δ::hphR* |
| TM252 | $h^{09}$ | *ura4-D18 leu1-32 swi6Δ::kamR* |
| TM886 | $h^{09}$ | *(Blp1)::LEU2 leu1::*$ura4^{+}$*-[mfm3p-YFP]-[map2p-CFP] ura4-D18 ade6-M216 swi6Δ::kamR shf1Δ::hphR* |
| PG1899 | *mat3-M(EcoRV)::ura4+* | *leu1-32 ura4-DS/E ade6-216* |
| PAE1900 | *mat3-M(EcoRV)::ura4+* | *leu1-32 ura4-DS/E ade6-216 swi6Δ::kamR* |
| PAE1901 | *mat3-M(EcoRV)::ura4+* | *leu1-32 ura4-DS/E ade6-216 dcr1Δ::hphR* |
| PAE1902 | *mat3-M(EcoRV)::ura4+* | *leu1-32 ura4-DS/E ade6-216 pcr1Δ::*$LEU2^{+}$ |
| PAE1903 | *mat3-M(EcoRV)::ura4+* | *leu1-32 ura4-DS/E ade6-216 dcr1Δ::hphR pcr1Δ::*$LEU2^{+}$ |
| PAE1904 | *mat3-M(EcoRV)::ura4+* | *leu1-32 ura4-DS/E ade6-216 shf1Δ::kamR* |
| PAE1905 | *mat3-M(EcoRV)::ura4+* | *leu1-32 ura4-DS/E ade6-216 dcr1Δ::hphR shf1Δ::kamR* |
| PAE1906 | *mat3-* | *leu1-32 ura4-DS/E ade6-216 pcr1Δ::*$LEU2^{+}$ *shf1Δ::kamR* |

| | | |
|---|---|---|
| | *M(EcoRV)::ura4+* | |
| PAE1907 | *mat3-M(EcoRV)::ura4+* | *leu1-32 ura4-DS/E ade6-216 set1Δ::kamR* |
| PAE1908 | *mat3-M(EcoRV)::ura4+* | *leu1-32 ura4-DS/E ade6-216 dcr1Δ::hphR set1Δ::kamR* |
| PAE1909 | *mat3-M(EcoRV)::ura4+* | *leu1-32 ura4-DS/E ade6-216 pcr1Δ::LEU2*$^{+}$ *set1Δ::kamR* |

### Supplementary Table S2. List of oligonucleotides used in this study

| Used for the gene deletion (used plasmid for amplification) | Oligonucleotide Sequence |
|---|---|
| shf1_ Forward (pFA6a- kanMX6 or -hphMX6) | 5'- ACCAATTGTTTTAATTAGATAGTGTACATTTTTCGTGAAAAA AAAGAACCATTAATATCAGCACTTAATTATTCACTTTAGCGC CAGATCTGTTTAGCTT-3' |
| shf1_Reverse (pFA6a- kanMX6 or -hphMX6) | 5'- AACTTTCAATAAAAAGTCATAATAATATCCACAAAGCGGAA GCTCAATGATTTGTACGCAGACGATGATTGCAATAATTATCG ATGAATTCGAGCTCGT-3' |
| set1_ Forward (pFA6a- kanMX6 or -hphMX6) | 5'- ATTTAGTGCACAGGTTTTAATATCTCTTGATTTTTTTTTTTT TTACTTTATAGCTCTTTTTTGTTTTGATCGAATGCCTGGGATT TGCTACTTGGATTGCGCCAGATCTGTTTAGCTT-3' |
| set1_ Reverse (pFA6a- kanMX6 or -hphMX6) | 5'- TTAAAAAATAAAAAGCATTAGTGTACCATCAGATATAATGC GTGCTTTTTAAACGAACTATTATAATTGTACAGCTGCCCATA TATTCATGTACCATCAAATCGATGAATTCGAGCTCGT-3' |
| rhp6_ Forward (pFA6a- kanMX6) | 5'- GGTGTAATTCCAAGGCGATATCGATATTTGTGCAACTTTTTT TTAAAGTTATCACAAATAGAAGAGAGGTTGCTATAAAAGGC GCGCCAGATCTGTTTAG-3' |
| rhp6_ Reverse (pFA6a- kanMX6) | 5'- AGATTTAATGTGAAAGGCGGTTGAAAAAGAAGAGTAAGTTC TAATGAAATAGGGATTATTAAGACAGCTATGTCTTGAAACA TCGATGAATTCGAGCTCG-3' |
| dcr1_ Forward (pFA6a -hphMX6) | 5'- AATAGCTTAGGATTCATTATTTTTAAGAGACAAATTTCTCG TCAATTGAATGAAACCTTCCGCCTTTATTTTCTTTTTGCGGAT CCCCGGGTTAATTAA-3' |
| dcr1_ Reverse (pFA6a -hphMX6) | 5'- GGAGACCCAAATTGAAAGTTTGAAAAGTTACAAGGGCCGCG GTCATAAAAAATGAAATACTGTATATTTCAAGTCTCAAGGA ATTCGAGCTCGTTTAAAC-3' |
| pcr1_ Forward (pREP1(LEU2)) | 5'- TCAATTTCCCATCTCCCCCTTTGTTTCCTTTGTTATATTTTTGT TTGTGATCATCTGATTCCCCCTTTTCTATACATTGACCTTATC ACGTTGAGCCATT-3' |
| pcr1_ Reverse (pREP1(LEU2)) | 5'- AGAACATTAGGCTAAAATAGGAGCCATAAATCTACATGCAA CATCACATTAAAATCAATTGCTTTATTGTAAAGGGGGGTCTA AGGCGCCTGATTCAAGA-3' |
| hht3_ Forward (pFA6a -hphMX6) | 5'- TAGGGACATATTTATACAGCGTAATCTCCCTATAATTTGTTT TTATCTACACAGGGCAATTATCAGAACACTAGAAAATACGG ATCCCCGGGTTAATTAA-3' |

| | |
|---|---|
| hht3_ Reverse (pFA6a -hphMX6) | 5'-GGGAAGGATTACAATGGGATAGACGACCAAAAGATTCCAAATCACACCATCAGATGGCAACCACAATTTGGTAAAGTTGCGAATTCGAGCTCGTTTAAAC-3' |
| hht1 and 2 gRNA_ Forward | 5'-TCTACCGGTGGTAAGGCACC-3' |
| hht1 and hht2 gRNA_ Reverse | 5'-GGTGCCTTACCACCGGTAGA-3' |
| Set1- Y897A gRNA_ Forward | 5'-TACTTATGATTACAAGTTTC-3' |
| Set1- Y897A gRNA_ Reverse | 5'-GAAACTTGTAATCATAAGTA-3' |
| **Multiplex PCR** | |
| FAM-MT1 | 5'-AAATAGTGGGTTAGCCGTGAAAGG-3' |
| MP1 | 5'-ATCTATCAGGAGATTGGGCAGGTG-3' |
| MM1 | 5'-GGGAACCCG CTGATAATTCTTGG-3' |
| **qPCR** | |
| *SRE2*_ Forward | 5'-ACCTTGTGGTTGATTTCACGTT-3' |
| *SRE2*_ Reverse | 5'-ACGGACTAACAAGGAAGCGT-3' |
| K_ Forward | 5'-GTTTGTAAGCGGCACATCCA-3' |
| K_ Reverse | 5'-AATCTTGCCTCCATCGTCCA-3' |
| *SRE3*_ Forward | 5'-TGCCAACATAACGATATCATCA-3' |
| *SRE3*_ Reverse | 5'-GCCTAGCGATTGATGTCAGTG-3' |
| *act1*_Forward | 5'-CTCAAAGCAAGCGTGGTATTT-3' |
| *act1*_ Reverse | 5'-TCTTTTCCATATCATCCCAGTTG-3' |

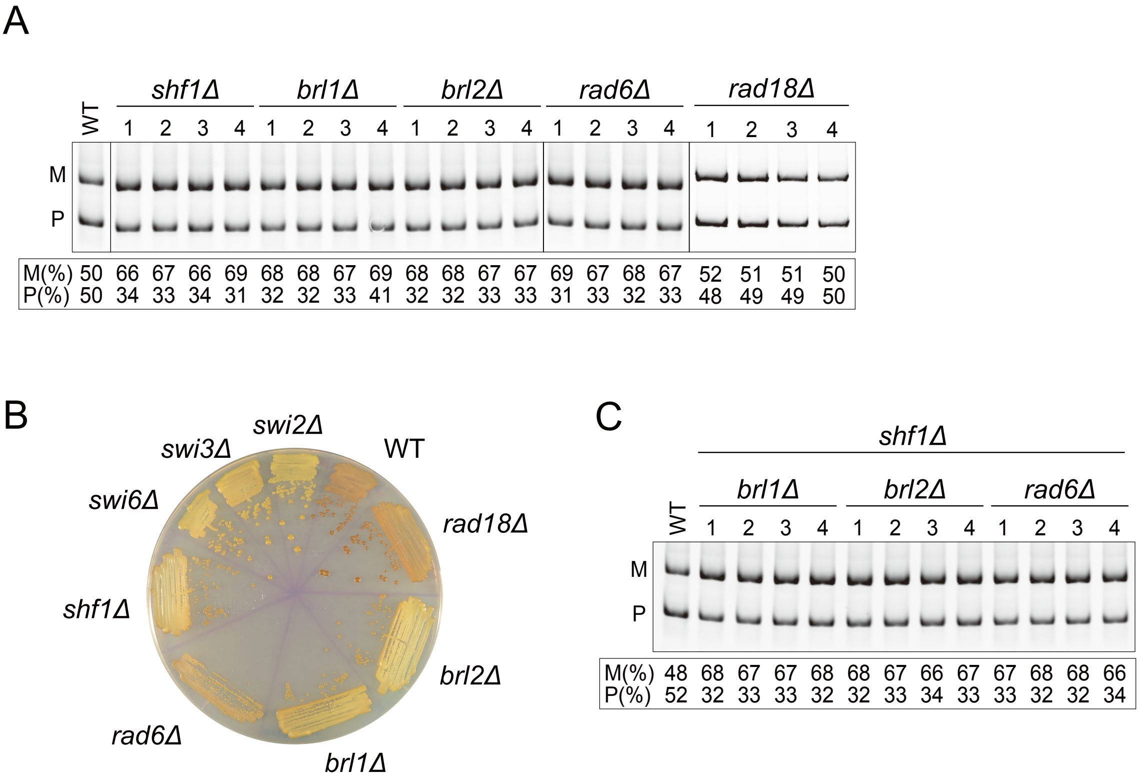


**Supplementary Fig. S1. Multiplex PCR analysis of single and double mutants in HULC subunits.**

(A, C) Gel pictures for the multiplex PCR analysis of the deletion mutants shown in Fig. 2. The *mat1-P* and *mat1-M* band intensities were measured and the % of P or M band intensity was calculated as P/(P+M)×100 or M/(P+M)×100 for each single deletion mutant of the subunits of HULC (A) or the double mutants (C). (B) Iodine staining of the single deletion mutant of HULC subunits and the *rad18*Δ mutant related to Fig. 2B.

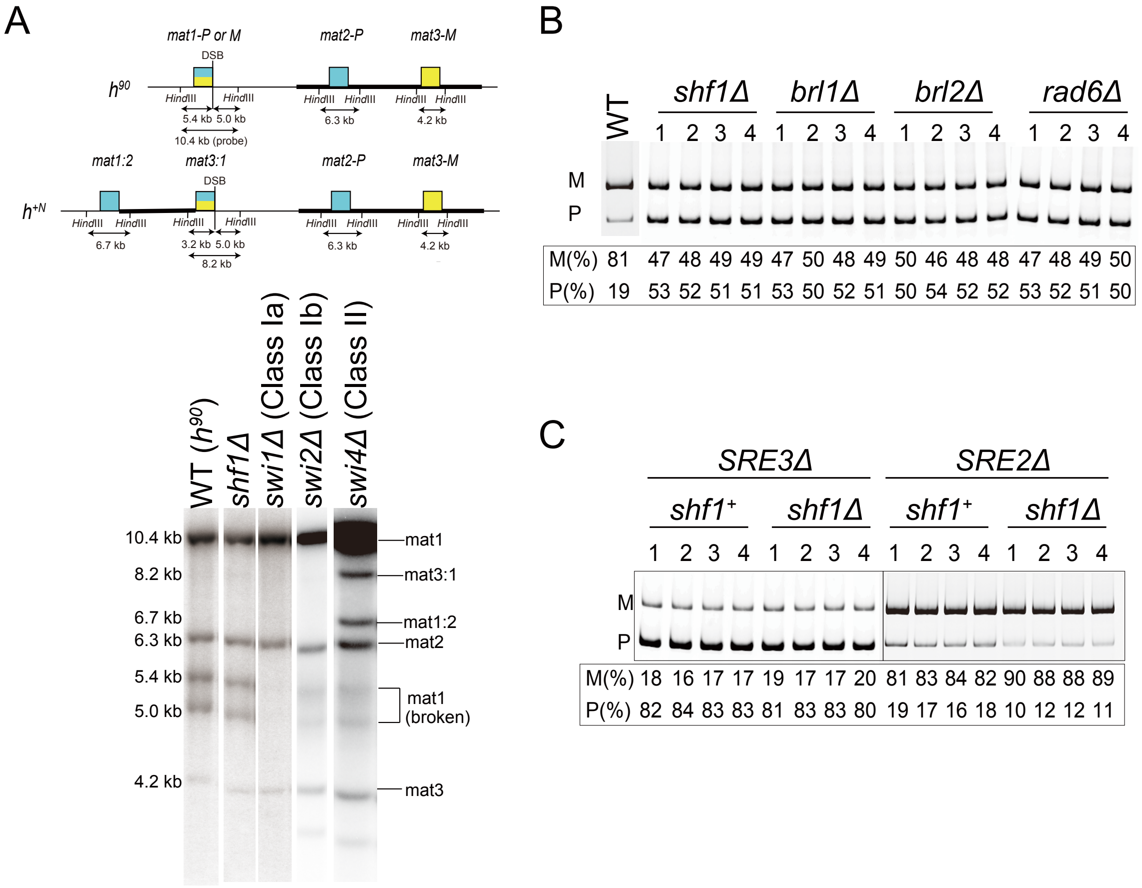


**Supplementary Fig. S2. Classification of *shf1*Δ MTS defect by Southern blot, and multiplex PCR analysis of HULC subunit mutations in $h^{09}$ background or in the presence of mutated *SRE* elements.**

(A) Southern blot of *Hin*dIII-digested genomic DNA. The probe was made from a 10.4 kb *mat1-P Hin*dIII fragment. The 5.4 kb and 5.0 kb fragments are cleavage products of the 10.4 kb *mat1* fragment, due to a double-strand break (DSB) generated by the imprinting at *mat1*. These fragments are absent in Class Ia mutants such as *swi1*Δ and *swi3*Δ that lack the imprint. Class Ib mutants as *swi2*Δ present the same band pattern as the $h^{90}$ WT strain. An 8.2 kb (*mat3:1*) and 6.7 kb (*mat1:2*) band would result from the $h^{+N}$ rearrangement of the *mat* region, which is detected in Class II mutants, but not seen in any of the mutants here. (B-C) Gel pictures for the multiplex PCR analysis of the deletion mutants shown in Fig. 3. The % of P or M band intensity was calculated as in Supplementary Fig. S1. (B) Single deletion mutants lacking subunits of HULC in $h^{09}$ background, shown in Fig. 3A. (C) Deletion of *SRE3* in *shf1*$^{+}$ or *shf1*Δ background and deletion of *SRE2* in *shf1*$^{+}$ or *shf1*Δ background, shown in Fig. 3B.

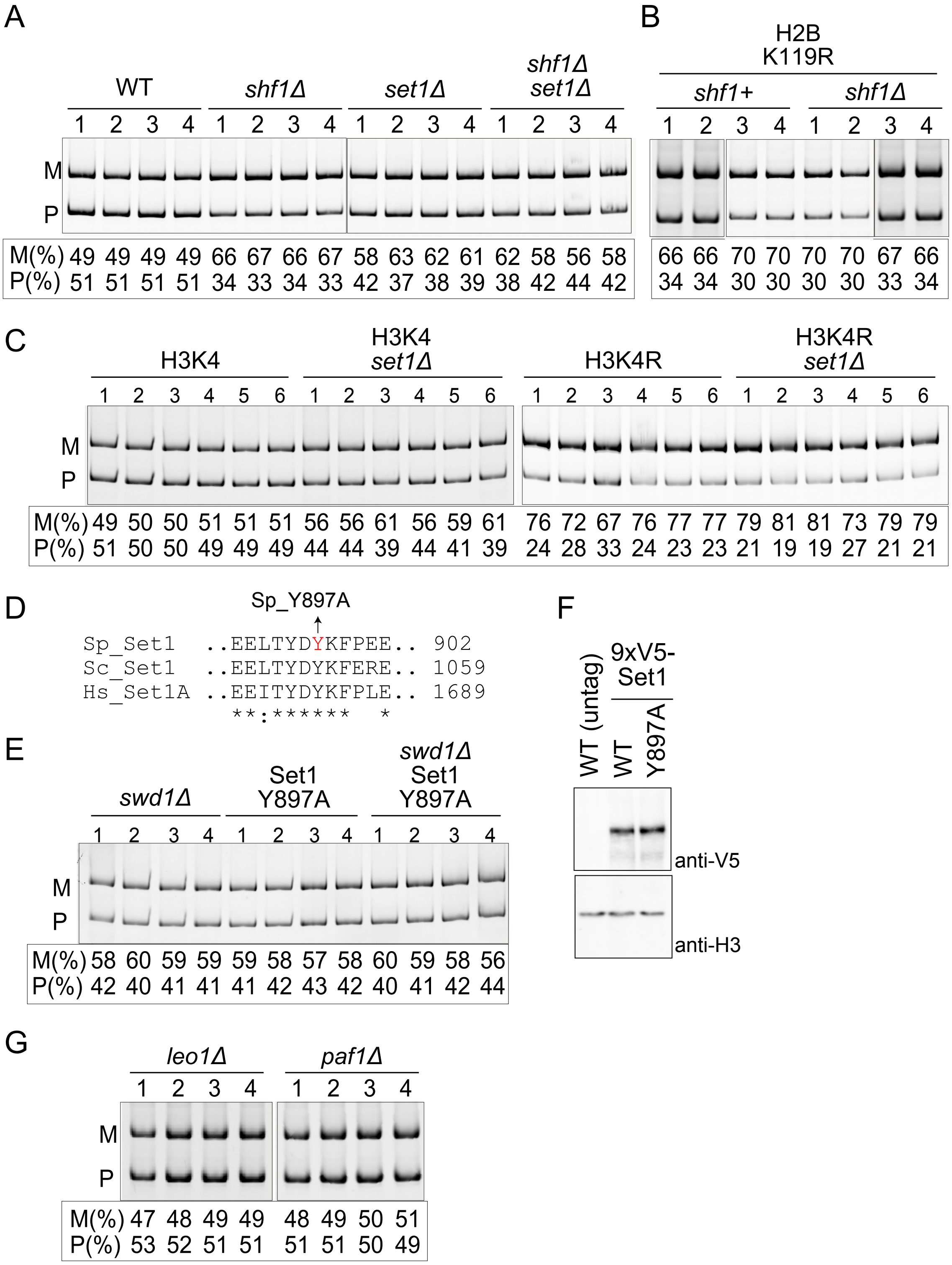


**Supplementary Fig. S3. Multiplex PCR analysis of mutants with defects in histones or various euchromatic factors.**

(A-C, E, G) Gel pictures for multiplex PCR analysis of the deletion mutants shown in Fig. 4. The % of P or M band intensity was calculated as in Supplementary Fig. S1 for (A) *shf1*Δ, *set1*Δ and *shf1*Δ *set1*Δ mutants, shown in Fig. 4A; (B) H2BK119R mutant in *shf1*$^{+}$ or *shf1*Δ background, shown in Fig. 4B; and (C) H3K4R mutant in *set1*$^{+}$ or *set1*Δ background, and catalytically dead Set1, shown in Fig. 4D. (D) Sequence alignment of the catalytic region of Set1. Sp: *S. pombe*, Sc: *S. cerevisiae,* Hs: *Homo sapiens*. (E) Multiplex PCR analysis of *swd1*Δ and Set1-Y897A single and double mutants. (F) Immunoblot with untagged, 9×V5-Set1 and 9×V5-Set1-Y897A strains. The primary antibodies used were anti-V5 or histone H3 as indicated. (G) Multiplex PCR analysis of Paf1C subunit deletion mutants, *leo1*Δ and *paf1*Δ*,* shown in Fig. 4E.

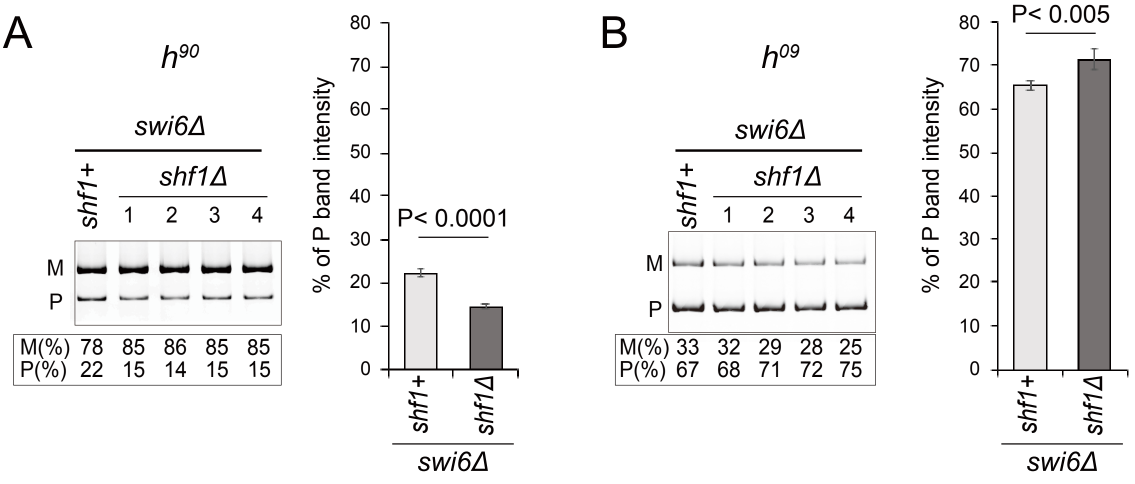


**Supplementary Fig. S4. Effects of *swi6* deletion on MTS in $h^{90}$ and $h^{09}$ cells with *shf1*$^{+}$ and *shf1*Δ backgrounds.**

Gel picture for multiplex PCR analysis and % of M and P band intensities calculated from: (A) $h^{90}$ strains with a *swi6* deletion in *shf1*$^{+}$ or *shf1*Δ background. A *swi6* deletion causes bias towards M cells which is slightly increased on the *swi6Δshf1Δ* double mutant. (B) $h^{09}$ strains, where the contents of the silent cassettes are swapped to *mat2-M mat3-P,* with a *swi6* deletion in *shf1*$^{+}$ or *shf1*Δ background. A *swi6* deletion causes bias towards P cells on $h^{09}$ cells, and a double mutant of *swi6Δshf1Δ* slightly increases this bias.

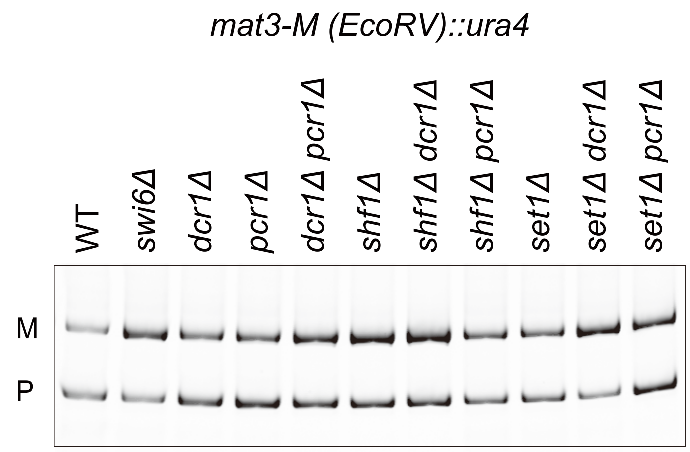


| | | | | | | | | | | | |
|---|---|---|---|---|---|---|---|---|---|---|---|
| M(%) | 41 | 68 | 47 | 46 | 56 | 61 | 61 | 66 | 53 | 74 | 50 |
| P(%) | 59 | 32 | 53 | 54 | 44 | 39 | 39 | 44 | 45 | 26 | 50 |

**Supplementary Fig. S5. Multiplex PCR analysis of strains used for the silencing assay.** Gel picture for multiplex PCR analysis and % of P and M band intensities for the strains used for the silencing assay in Fig. 6C.

1 AGCGCCCAAT ACGCAAACCG CCTCTCCCCG CGCGTTGGCC GATTCATTAA
51 TGCAGCTGGC ACGACAGGTT TCCCGACTGG AAAGCGGGCA GTGAGCGCAA
101 CGCAATTAAT GTGAGTTAGC TCACTCATTA GGCACCCCAG GCTTTACACT
151 TTATGCTTCC GGCTCGTATG TTGTGTGGAA TTGTGAGCGG ATAACAATTT
201 CACACAGGAA ACAGCTATGA CCATGATTAC GCCAAGCTTG TCGATCGACT
251 ACGTCGTTAA GGCCGTTTCT GACAGAGTAA AATTCTTGAG GGAACTTTCA
301 CCATTATGGG AAATGGTTCA AGAAGGTATT GACTTAAACT CCATCAAATG
351 GTCAGGTCAT TGAGTGTTTT TTATTTGTTG TATTTTTTTT TTTTTAGAGA
401 AAATCCTCCA ATATATAAAT TAGGAATCAT AGTTTCATGA TTTTCTGTTA
451 CACCTAACTT TTTGTGTGGT GCCCTCCTCC TTGTCAATAT TAATGTTAAA
501 GTGCAATTCT TTTTCCTTAT CACGTTGAGC CATTAGTATC AATTTGCTTA
551 CCTGTATTCC TTTACATCCT CCTTTTTCTC CTTCTTGATA AATGTATGTA
601 GATTGCGTAT ATAGTTTCGT CTACCCTATG AACATATTCC ATTTTGTAAT
651 TTCGTGTCGT TTCTATTATG AATTTCATTT ATAAAGTTTA TGTACAAATA
701 TCATAAAAAA AGAGAATCTT TTTAAGCAAG GATTTTCTTA ACTTCTTCGG
751 CGACAGCATC ACCGACTTCC GTGGTACTGT TGGAACCACC TAAATCACCA
801 GTTCTGATAC CTGCATCCAA AACCTTTTTA ACTGCATCTT CAATGGCCTT
851 ACCTTCTTCA GGCAAGTTCA ATGACAATTT CAACATCATT GCAGCAGACA
901 AGATAGTGGC GATAGGGTTG ACCTTATTCT TTGGCAAATC TGGAGCAGAA
951 CCGTGGCATG GTTCGTACAA ACCAAATGCG GTGTTCTTGT CTGGCAAAGA
1001 GGCCAAGGAC GCAGATGGCA ACAAACCCAA GGAACCTGGG ATAACGGAGG
1051 CTTCATCGGA GATGATATCA CCTGCATCTT AGGTTAATTA TTAATTAATC
1101 TCATCGATCG CTTTCTTCTT ACTTTAACAA AAACACATCG AAAGAGATAC
1151 GATGTTTGAT TTGAATACTT GTCAACGTAA AAATATCTTA GAACTTCAGC
1201 CTTATCGCTG TGCGAGAGAG TATGTAATAA CTTGATAAAC TAGAAAGCGT
1251 GATATATATT AACATAGAAT ATAGTGACTT CTCTGAGTAT GTTTTATCTT
1301 TATATTTGGG TAATAATTGA TATGAGGGCT ACTTCTAGAA AGATATTTTA
1351 TTAGCTGCTT GGAAAGCAAT GAAGTACTCC CTGTTTACTG TTGTTGTTGG
1401 AGCTGGCGAT TTCAGGCCTT TGTTGTTTTT TACCCTTTTC TCGCCTATAT
1451 ATTTTTTGCT AATCTTAATA GAGGTGTCCT TCTTGATGCC AATGAATGTG
1501 CTTATGGTTC GGTTATTTCC GTGGACGGTG TGGAATTCAA TCGTTATCCA
1551 GACCCTAGAC AGATCGAAGT AAAGCAAAGA TTATGTGATT TGAGAAACAA
1601 AGAGCTTTCC ATTACCAAGC CACTAACACC AGATAACATT TGCATGGGTG
1651 TTGGCAGCGA TGAAATTATT GACTCCTTAA TTCGCATTTC TTGTATTCCT
1701 GGAAAAGATA AGATTTTGAT GTGTCCACCT TCGTACGGAA TGTATACGGT
1751 CTCTGCAAAA ATCAACGATG TTGAGGTTGT TAAGGTTCTC CTTGAACCAG
1801 ATTTCAATTT GAACGTCGAT GCTATTTGCG AAACACTTTC TAAAGATAGT
1851 GCCATCAAAG TCTTTTTTGC TTGTTCCCCT GGTAATCCAA CAGCTAAGGC
1901 TCTTAAACTA GAAGATATTA AGAAGATTTT GGAGCACCCT ACATGGAATG
1951 GAATCGTTGT CGTCGATGAG GCCTATATTG ATTTCTCTGC TCCCGATATG
2001 TCTGCCTTAA CTCTTGTCAA TGAATATCCA AACCTTGCTG TTTGCCAAAC
2051 CCTTTCGAAA TCTTTCGGTC TAGCAGGAAT TCGGTAAGTT AATCTAAGCT
2101 TTTAAGCATT AATATTATTT CTTTACTAAC TATGATGTTT AGTATCGGAT
2151 TTTGTTTGAC AAGTAAACCT ATTGCTACCA TTATGAACTC ATTAAAGGCT
2201 CCTTATAACA TTAGCGAACC CACCTCTCGT TTAGCTTTGG ATGCTTTATC
2251 CCCTCAGTCG ATTGATAAAA TGCATACATA TAGAGATGCT ATTATACAGC
2301 AAAGAGTTCG TCTTTGCAAA GAGCTTACTA CCATTAAAGG AATGGGCAAG
2351 ATTATAGGTG GTTATGATGC AAATTTTATC CTCATTCAGG TTTTAGATAG
2401 ACCAGAAGGA GGTAAGCCTA GTAACGATGC CGCAAAATAT CTTTATTTAC
2451 AAATGGCCAC GATGCACAAA GTCGTTGTTA GATTTAGAGG TACAGAGCCT
2501 CTTTGCGAAG GAGCATTGCG TATCACTGTT GGCACAGAAG AAGAGAATAC
2551 GATATTACTA AAAACTATTA AACTTGTATT GCAGGAATAC TATACAAAAA

2601 AATGAGGTTT CCTTTTCTGG CCTATTAGAA CGTTTCAATT AGACAAATTA
2651 TAACTCTTTC CTTTCCCCTT CCGGACTCGT CACATTTCTT TTCTGATGTA
2701 CCCAATATAT TTTGTGTTCG CTAATTGCGC TTGCATTCCT TCTTTCTTTA
2751 TTAAAATACG TTTTCTTCCT TTTTGGTGTG TTTCCGTATA GTTCGATAAG
2801 CAATGAAAAG ATGTTATTTT TAATTTTGTA AAGGAGTTAC GCTAATTTAT
2851 CGAATTCCTG CAGGTCGATC GACTCTAGAG GATCAGAAAA TTATCGCCAT
2901 AAAAGACAGA ATAAGTCATC AGCGGTTGTT TCATTTCCTA TATTTTTTTT
2951 TTATTTTTTT ATTTTTTAAT AAGGGAAAAT TTAACGTCTA AGGATACAGA
3001 AGATTGTTAG CACATTAAAG TAATAAAGGC TTAAGTAGTA AGTGCCTTAG
3051 CATGTTATTG TATTTCAAAG GACATAATCT AAAATAATAA CAATATCATT
3101 TCTCACAAGT TATTCAATTT TCTTTTTTTT TTCTAATAAT ATCAAGAATG
3151 TATTATTTGT TTGACATAAG TCAACTAATT TATTTAATAT GCTGGATTAA
3201 TCTTGCAGAC ATGTAAATTA ACAAGTTTTA GTCAAATAAC GTTGAAGTTT
3251 CAATGAACTC AAATAATTTC TCTTTTTTTT TATATAACCA TATGTCTAAT
3301 CTGATTTATA TTTTCCGCAG GGATCAACTG AAGTTATGAC ATTTGGATTG
3351 GATCACTTAT AACCTTGGTC GCCAAATAAT ACAAAAATCA GCGTTATAAA
3401 ACAAAGAAGG TTTTTGTTAA GAAATTAATC CTCTTTCTTG ATAAGAAAGT
3451 TGAACCGAAA TTGCAGATAC TGATATATGA AAATAATACC CACAATTTTG
3501 GGAATAGCGC AAGCCTCAAT TTAAACAATA GGTGAGGACA CATGATAATG
3551 ACCTCAATGA TTGTTAGAAG AAAAGAGCCT CATTACAAAA TCGAAAAATG
3601 AATGGTTGGG TACAAGTTTC CAAAACATGG TAAAGTGGAC TTTGCGTATG
3651 AGACGTAAAT AGAAAAAAAC ACTTGTTATA TGTTTTCTAG AATTATTGTT
3701 GTCTCTTTAT GGTTGGATGA TGCAAAATAG TAATTTCGGT TAGTTGCTGT
3751 AAAACACCAC GAGACAAATA GATATGGATA TTTATTAAAT CAGGAAAAAC
3801 GTAACTCTCG GCTACTGGAT GGTTCAGTCA CCCAACGATT ACTGGGGAGA
3851 GAAAACAGGG CAAAAGCAAA GCTTAAAGGA ATCCGATTGT CATTCGGCAA
3901 TGTGCAGCGA AACTAAAAAC CGGATAATGG ACCTGTTAAT CGAAACATTG
3951 AAGATATATA AAGGAAGAGG AATCCTGGCA TATCATCAAT TGAATAAGTT
4001 GAATTAATTA TTTCAATCTC ATTCTCACTT TCTGACTTAT AGTCGCTTTG
4051 TTAAATTGGC CTCGTTTGGC CTGATGAGTC CGTGAGGACG AAACGAGTAA
4101 GCTCGTCGCC AAACAAATTG ATTGGTTTTA GAGCTAGAAA TAGCAAGTTA
4151 AAATAAGGCT AGTCCGTTAT CAACTTTAAA AAGTGGCACC GAGTCGGTGC
4201 TTTTAGATAA GTCACTATGT CCGAGTGGTT AAGGAGTTAG ACTCGAATTC
4251 CTACATTCGT GGCATCTAAT GGGCTCTGCC CGCGCAGGTT CAAATCCTGC
4301 TGGTGACGGG CGTCTTCACT AGAAGACGCG TTTTAGAGCT AGAAATAGCA
4351 AGTTAAAATA AGGCTAGTCC GTTATCAACT TGAAAAAGTG GCACCGAGTC
4401 GGTGCTTTTG GCCGGCATGG TCCCAGCCTC CTCGCTGGCG CCGGCTGGGC
4451 AACATGCTTC GGCATGGCGA ATGGGACCCG GGTAAAAGGA ATGTCTCCCT
4501 TGCCAGTACT GCTAGGGTTT TTCTTTCAAA CTATGGAAGC CCATTCAAGC
4551 TGCATATTAC GATTTTGTTT TTCGCTTTTA GAAAGTGGTT TAGATGAGAT
4601 AATAGAAAAA TTCTTGATCT CCGACAACGA GTACTTTTAT TTTTTTTGCT
4651 AATCACTTTA CTCAATATTA GCTCGAAATC GTAGAAACGT AGACGGGTGC
4701 GGGATACCGA GTGGTGTAGT TAAGAATTTT TATAAACCAC GTGGCCCAAA
4751 AATATGAACC CAAAACGTTT ATACATGAGT ATACTTTAAG AAGGCTATAC
4801 CCCTTCGTGT TAGATGTAGT TTTAGCTACC CAACCCGAGT CTATGAGCTT
4851 GACTTCAGAT GTAGAAGGCA TTAAATCGTT TTGAATATTA ATTAAAAAAC
4901 GATGAAAATT AAATATTTAA AAGCAATCAT ACGCTGAAAA TTTAGTGCTG
4951 TGGCTAATCC TTCAACATGG AAATGCCATA AAAGTGACTT TGACAAAAAA
5001 AAAAGTATAT ACAGGTAGTA AACTCATCTA CTTCATTGAC TTTGTTTACA
5051 GCATGTGGAA GGAGGAATAT TTATTGCTAA ATCGTAGTTT AACATTCAAT
5101 AAGTAATACT ATTGAAATTC GACAAGATTG GCCGCATGGA TGAAAAAGAG
5151 GCATTTTGCT TTGGGAGAAT TAGTTCAAAT TAGAACTGAA AAAAAAAACT

5201 TTACGAGGCA AAAATGTCGG ATTGAGATCG TAAAAGTTCG CTCGTCGTCT
5251 TTTGCTTTGT GATTGTTTTC ATGGATACAT CTTGCTGGAT ATTTAAATTT
5301 TAGTACTATG TATAAGATAT TCTATAAATG TTTTATCACC CAAACCTGTT
5351 AGCGCCTTCT TAATTCTATT CAATCTGGCT TTTGCTCTGA GACTACTTCT
5401 TGGACTTTCA CTACTTGTTA GTTATACGGA ATTTGTGTAA TTAGAAGTGA
5451 AATAATCCTT TCTATTAGTA ATGCGAGCTC GAATTCGAGT CTAACTCCTT
5501 AACCACTCGG ACATAGTGAC TTATCTGACA CTCATTGTAA ATTAATATAT
5551 ATATAGGCAT TTTGTTTAGT TAAAGGTACT TAAGTAATTA GTATAAACGA
5601 ACCAATTTTA TAATCAGGAA GTTAAGTGAA TGGTAGCACA TGTCGTAAAA
5651 ATTGTGAATT TTTATTGAAT AATATTTTAA ATACAAGCCT TTCTAGACTA
5701 GGTATACTCA TAAACATATA TGAGCAAAAG GATAGAGGAG ATTACATTGC
5751 ATCTTCTACA AATTTATTTA TTGCCCTTTA CTGAAAAATT AAATAATGAG
5801 TACTAAATGA TAAAAAGCGC TCAGTACAAG AAAGATTGCA AAAATATTGC
5851 ATTCTTCATG AATTAAAGTT GCATATAAGG CATATTGAAA GTAATAGTAC
5901 TAAAACAGCA GTTAGCGAAA ATTAATAGAA TTATATTCGC AAGACAATTG
5951 TGACATTAAA TTAAAAAATT GTAGAATTTT TACTATCCTC TTTAACGCCA
6001 TGAGCCTTTA TAAAAAGGTT AAATTAGTTT TAACATTCTT TTTTTGAGTA
6051 AGAGTTTACA ATTTATCAAA ACCTGTGTTA TTATATTCAT TAGTTTCAAT
6101 TTATTAGCAT CTAGAGAAAA ATCAATTGGC AGTTACATTG TTGGAATTTA
6151 TGAAGAAAAA GAATCTACAA CGGAGAATAA GTTGCTGATC GCTTTCCCTA
6201 AAATTGTATA TTTTGCTGAG CTTATTTTGA CATTTCGTTG AAGTTTTCTC
6251 TAATTCGCAT TCATTTTAAG TAAAACAATG AGAAATAAAA TTACAAAAAA
6301 TACAAATTAA AATACAATTT TTAGCTATAA TTATAGACGA TGCCCTTGTA
6351 TCCCATTCTG TCTCGCTTGC CCCTACTTTT TATCTTTTAT ATACCATAAT
6401 GAACGCTGCC GCTACTAACC ATACCCCGAT TTTACATTTC GGACTCCCAA
6451 GGACGTACAA AATAGAAAAC TATAGAAAAA AATAATCAGA AAATAGCATG
6501 TCATCTCTTT GTAAAACGCG TTTGCAAGAA GAAAGGAAAC AATGGAGAAG
6551 AGATCATCCA TTTGTATGTA AATTTTAGTA AACTTGAAGA AATCACTAAC
6601 AACTTCTCTT ACTTAGGGAT TCTATGCAAA ACCTTGTAAA TCATCTGATG
6651 GAGGACTCGA TTTAATGAAT TGGAAGGTTG GAATTCACTG GCCGTCGTTT
6701 TACAACGTCG TGACTGGGAA AACCCTGGCG TTACCCAACT TAATCGCCTT
6751 GCAGCACATC CCCCTTTCGC CAGCTGGCGT AATAGCGAAG AGGCCCGCAC
6801 CGATCGCCCT TCCCAACAGT TGCGCAGCCT GAATGGCGAA TGGCGCCTGA
6851 TGCGGTATTT TCTCCTTACG CATCTGTGCG GTATTTCACA CCGCATACGT
6901 CAAAGCAACC ATAGTACGCG CCCTGTAGCG GCGCATTAAG CGCGGCGGGT
6951 GTGGTGGTTA CGCGCAGCGT GACCGCTACA CTTGCCAGCG CCCTAGCGCC
7001 CGCTCCTTTC GCTTTCTTCC CTTCCTTTCT CGCCACGTTC GCCGGCTTTC
7051 CCCGTCAAGC TCTAAATCGG GGGCTCCCTT TAGGGTTCCG ATTTAGTGCT
7101 TTACGGCACC TCGACCCCAA AAAACTTGAT TTGGGTGATG GTTCACGTAG
7151 TGGGCCATCG CCCTGATAGA CGGTTTTTCG CCCTTTGACG TTGGAGTCCA
7201 CGTTCTTTAA TAGTGGACTC TTGTTCCAAA CTGGAACAAC ACTCAACCCT
7251 ATCTCGGGCT ATTCTTTTGA TTTATAAGGG ATTTTGCCGA TTTCGGCCTA
7301 TTGGTTAAAA AATGAGCTGA TTTAACAAAA ATTTAACGCG AATTTTAACA
7351 AAATATTAAC GTTTACAATT TTATGGTGCA CTCTCAGTAC AATCTGCTCT
7401 GATGCCGCAT AGTTAAGCCA GCCCCGACAC CCGCCAACAC CCGCTGACGC
7451 GCCCTGACGG GCTTGTCTGC TCCCGGCATC CGCTTACAGA CAAGCTGTGA
7501 CCGTCTCCGG GAGCTGCATG TGTCAGAGGT TTTCACCGTC ATCACCGAAA
7551 CGCGCGAGAC GAAAGGGCCT CGTGATACGC CTATTTTTAT AGGTTAATGT
7601 CATGATAATA ATGGTTTCTT AGACGTCAGG TGGCACTTTT CGGGGAAATG
7651 TGCGCGGAAC CCCTATTTGT TTATTTTTCT AAATACATTC AAATATGTAT
7701 CCGCTCATGA GACAATAACC CTGATAAATG CTTCAATAAT ATTGAAAAAG
7751 GAAGAGTATG AGTATTCAAC ATTTCCGTGT CGCCCTTATT CCCTTTTTTG

7801 CGGCATTTTG CCTTCCTGTT TTTGCTCACC CAGAAACGCT GGTGAAAGTA
7851 AAAGATGCTG AAGATCAGTT GGGTGCACGA GTGGGTTACA TCGAACTGGA
7901 TCTCAACAGC GGTAAGATCC TTGAGAGTTT TCGCCCCGAA GAACGTTTTC
7951 CAATGATGAG CACTTTTAAA GTTCTGCTAT GTGGCGCGGT ATTATCCCGT
8001 ATTGACGCCG GGCAAGAGCA ACTCGGTCGC CGCATACACT ATTCTCAGAA
8051 TGACTTGGTT GAGTACTCAC CAGTCACAGA AAAGCATCTT ACGGATGGCA
8101 TGACAGTAAG AGAATTATGC AGTGCTGCCA TAACCATGAG TGATAACACT
8151 GCGGCCAACT TACTTCTGAC AACGATCGGA GGACCGAAGG AGCTAACCGC
8201 TTTTTTGCAC AACATGGGGG ATCATGTAAC TCGCCTTGAT CGTTGGGAAC
8251 CGGAGCTGAA TGAAGCCATA CCAAACGACG AGCGTGACAC CACGATGCCT
8301 GTAGCAATGG CAACAACGTT GCGCAAACTA TTAACTGGCG AACTACTTAC
8351 TCTAGCTTCC CGGCAACAAT TAATAGACTG GATGGAGGCG GATAAAGTTG
8401 CAGGACCACT TCTGCGCTCG GCCCTTCCGG CTGGCTGGTT TATTGCTGAT
8451 AAATCTGGAG CCGGTGAGCG TGGGTCTCGC GGTATCATTG CAGCACTGGG
8501 GCCAGATGGT AAGCCCTCCC GTATCGTAGT TATCTACACG ACGGGGAGTC
8551 AGGCAACTAT GGATGAACGA AATAGACAGA TCGCTGAGAT AGGTGCCTCA
8601 CTGATTAAGC ATTGGTAACT GTCAGACCAA GTTTACTCAT ATATACTTTA
8651 GATTGATTTA AAACTTCATT TTTAATTTAA AAGGATCTAG GTGAAGATCC
8701 TTTTTGATAA TCTCATGACC AAAATCCCTT AACGTGAGTT TTCGTTCCAC
8751 TGAGCGTCAG ACCCCGTAGA AAAGATCAAA GGATCTTCTT GAGATCCTTT
8801 TTTTCTGCGC GTAATCTGCT GCTTGCAAAC AAAAAAACCA CCGCTACCAG
8851 CGGTGGTTTG TTTGCCGGAT CAAGAGCTAC CAACTCTTTT TCCGAAGGTA
8901 ACTGGCTTCA GCAGAGCGCA GATACCAAAT ACTGTCCTTC TAGTGTAGCC
8951 GTAGTTAGGC CACCACTTCA AGAACTCTGT AGCACCGCCT ACATACCTCG
9001 CTCTGCTAAT CCTGTTACCA GTGGCTGCTG CCAGTGGCGA TAAGTCGTGT
9051 CTTACCGGGT TGGACTCAAG ACGATAGTTA CCGGATAAGG CGCAGCGGTC
9101 GGGCTGAACG GGGGGTTCGT GCACACAGCC CAGCTTGGAG CGAACGACCT
9151 ACACCGAACT GAGATACCTA CAGCGTGAGC TATGAGAAAG CGCCACGCTT
9201 CCCGAAGGGA GAAAGGCGGA CAGGTATCCG GTAAGCGGCA GGGTCGGAAC
9251 AGGAGAGCGC ACGAGGGAGC TTCCAGGGGG AAACGCCTGG TATCTTTATA
9301 GTCCTGTCGG GTTTCGCCAC CTCTGACTTG AGCGTCGATT TTTGTGATGC
9351 TCGTCAGGGG GGCGGAGCCT ATGGAAAAAC GCCAGCAACG CGGCCTTTTT
9401 ACGGTTCCTG GCCTTTTGCT GGCCTTTTGC TCACATGTTC TTTCCTGCGT
9451 TATCCCCTGA TTCTGTGGAT AACCGTATTA CCGCCTTTGA GTGAGCTGAT
9501 ACCGCTCGCC GCAGCCGAAC GACCGAGCGC AGCGAGTCAG TGAGCGAGGA
9551 AGCGGAAG

Features

1064-2846/*his3*

2862-3870/*nmt*1 promoter

4064-4106 /HHR (hammer-head ribozyme)

4107-4206 /ade6-M375 gRNA

4207-4311 /$tRNA^{Ser}$

4308-4335 /gRNA-array insertion site (*Bbs*I bi-directional restriction site)

4332-4411 /gRNA scaffold

4412-4479 /HDV ribozyme

4479-5479 /*nmt*1 term

5481-6681 /*ars*1

7758-8618 /ampR

6918-7344 /f1 ori

8766-9433 /pUC ori

**Supplementary Fig. S6. Nucleotide sequence of plasmid pEM59, allowing expression of an inserted gRNA sequence.**

hht1-H3K4R (HR donor)
5′-
CTGCAGTACGCTTGCGTTTCCATTAATTCTAAAGATCAACAATTGGCAAAAGTAGCACAACAGC
TATTTTTTTCTCGATTGTCTTTTTATTTGATATTCATTCTACTAGCTTGATATAATGGCTCGTA
CTAGACAAACAGCTCGTAAGTCTACCGGCGGTAAGGCACCCCGTAAGCAATTGGCCTCTAAGGC
CGCTCGTAAGGCCGCTCCCGCTACCGGAGGTGTTAAGAAGCCTCATCGTTATCGTCC-3′

hht2-H3K4R (HR donor)
5′-
GGGAAGCCGAAATCGCAATCTTAAATCAGGGTTAGGGTTGTGATTGACTGAGGTATATATAGCA
GGAAGTGTCCACACCCGACGTGGAAAGAACCTTTTTGTAAGTTTATTTACCGAATTACGTTATG
GCTCGTACCAGGCAAACTGCTCGTAAATCTACCGGCGGTAAGGCACCCCGTAAGCAATTGGCCT
CTAAGGCTGCCCGTAAGGCCGCTCCCGCTACTGGCGGTGTCAAGAAGCCTCATCGTTATCGTCC
TGG-3′

ade6 (HR donor)
5′-
GTGGTCAATTGGGCCGTATGATGGTAGAGGCAGCCCATCGCTTAAACATCAAATGCATCATCTT
GGATGCAGCAAATTCTCCTGCCAAACAAATTGATGGAGGACGTGAGCACATTGATGCATCATTT
ACTGACCCCGATGCAATTGTTGAACTGTCTAAGAAGTGCACG-3′

set1-Y897A (HR donor)
5′-
GGGAAATATCGCGCGTTTCATCAATCATTCATGCGCTCCTAATTGTATAGCTAGGATTATTAGA
GTTGAAGGGAAAAGAAAAATCGTAATTTATGCTGACAGGGATATTATGCATGGAGAGGAACTTA
CTTATGATGCCAAGTTTCCGGAAGAAGCTGATAAGATTCCTTGTTTGTGTGGTGCTCCAACATG
TCGTGGCTATTTAAACTAG-3′

**Supplementary Fig. S7. Nucleotide sequence of donor template used for CRISPR/Cas9-mediated genome editing.**

### SUPPLEMENTAL MATERIALS AND METHODS

#### Southern blot.

*S. pombe* cells were propagated in liquid medium overnight at 30°C to saturation and genomic DNA for Southern blots was prepared as described in Maki et al., 2018. Genomic DNA was digested with *HindIII* to examine the structure of the mating-type region. The digested samples were electrophoresed in 0.7% agarose gels. The probe used was a 10.4 kb *HindIII* fragment containing *mat1-P*.

#### Western blotting.

Culture aliquots of ~$5.0 \times 10^8$ *S. pombe* cells were washed in 20% TCA, pelleted and stored at -80°C until ready for use. Aliquots were thawed, resuspended in 300 µl 20% TCA and disrupted by a multi-beads shocker (Yasui Kikai) using 0.71- 1.18 mm glass beads at 4°C, and 15 cycles of 1 min on and 1 min off. Samples were collected by centrifugation, then washed with 5% TCA. Precipitates were resuspended in 30 µl 1M Tris-HCl (pH 8.0) and 270 µl 1.5x SDS sample buffer was added. 20 µl of each sample was separated on SDS–polyacrylamide gels, transferred to PVDF membrane with a Biorad semidry transfer apparatus at 15 V for 45 min. Blots were incubated with anti-V5 tag mouse monoclonal antibody (FUJIFILM) or anti-histone H3 rabbit polyclonal antibody (ab18521, Abcam). After washing, blots were incubated with either HRP-conjugated anti-mouse or anti-rabbit secondary antibodies (Jackson Immunoresearch and Cytiva, respectively), followed by treatment with ImmunoStar Zeta chemiluminescence reagent (FUJIFILM). Western blots were imaged and quantified using LAS4000 with ImageQuant software (Cytiva).
